# Hemin availability induces coordinated DNA methylation and gene expression changes in *Porphyromonas gingivalis*

**DOI:** 10.1101/2022.03.14.484211

**Authors:** Ricardo Costeira, Joseph Aduse-Opoku, Jon J Vernon, Francisco Rodriguez-Algarra, Susan Joseph, Deirdre A Devine, Philip D Marsh, Vardhman Rakyan, Michael A Curtis, Jordana T Bell

## Abstract

Periodontal disease is a common chronic inflammatory disease. *Porphyromonas gingivalis* is an important bacterium in the development of the disease and expresses a variety of virulence determinants. Hemin (iron [III] protopotphyrin IX), an essential nutrient of this organism, whose concentration increases with increasing inflammation, is a global regulator of virulence in *P. gingivalis*: high hemin levels increase expression of several virulence determinants. However, the mechanism through which hemin influences bacterial gene expression is poorly understood. Bacterial DNA methylation has the potential to fulfil this mechanistic role. Here, we characterised the methylome of *P. gingivalis*, and compared its variation to transcriptomic changes in response to changes in hemin concentration.

Gene expression and DNA methylation profiling of *P. gingivalis* W50 was performed, following continuous culture in chemostats with excess or limited hemin, using Illumina RNA-Seq and Nanopore DNA sequencing. DNA methylation quantification was carried out for Dam/Dcm motifs and all-context N6-methyladenine (6mA) and 5-methylcytosine (5mC) base pair modifications. Differential expression and methylation in response to excess hemin availability are presented after multiple testing correction (FDR 5%).

In excess hemin there were 161 over- and 268 under-expressed genes compared to limited hemin. Genes under-expressed in excess hemin were involved in iron recruitment (the hemophore HmuY) and transport (TonB-dependent receptors), and those over-expressed were involved in iron-sulphur cluster binding. Hemin-dependent differentially methylation was observed for the Dam ‘GATC’ motif and all-context 6mA and 5mC, with 36, 49 and 47 signals, respectively. Coordinated genome-wide differential expression and methylation effects were observed in 6 genes encoding a Ppx/GppA family phosphatase, a lactate utilization protein, a 4-alpha-glucanotransferase, two ABC transporter proteins, and a hypothetical protein HMPREF1322_RS00730. The findings indicate that altered genome methylation occurs in response to the availability of hemin and give insights into the molecular mechanisms of regulation of virulence in this bacterium.

**Author Summary:** DNA methylation has important roles in bacteria, including in the regulation of transcription. *Porphyromonas gingivalis,* an oral pathogen in periodontitis, exhibits well-established gene expression changes in response to hemin availability. However, the gene regulatory processes underlying these effects remain unknown. To this end, we profiled the novel *P. gingivalis* epigenome, and assessed epigenetic and transcriptome variation under limited and excess hemin conditions. As expected, multiple gene expression changes were detected in response to limited and excess hemin conditions that reflect conditions associated with health and disease, respectively. Notably, we also detected differential DNA methylation signatures for the Dam ‘GATC’ motif and both all-context N6-methyladenine (6mA) and 5-methylcytosine (5mC) in response to hemin availability. Joint analyses identified a subset of coordinated changes in gene expression, 6mA, and 5mC methylation that target genes involved in lactate utilization and ABC transporters. The results identify novel regulatory processes underlying the mechanism of hemin regulated gene expression in *P. gingivalis*, with phenotypic impacts on its virulence in periodontal disease.

## Introduction

Teeth are secured in the mouth through the periodontal ligament, which is attached to the subgingival root surface and the alveolar bone. Periodontal disease is a common chronic inflammatory disease of the periodontal tissues that is driven by bacteria in subgingival microbial biofilms on the root surface. In periodontal disease the integrity of the attachment tissues becomes compromised leading to bleeding, gingival recession, formation of deep periodontal pockets and periodontal bone loss; ultimately culminating in tooth loss in a susceptible host. Periodontitis is a global health issue affecting both developed and developing nations (1, 2). Furthermore, periodontitis is also associated with a range of extra-oral inflammatory conditions, such as diabetes and cardiovascular disease, rheumatoid arthritis, Alzheimer’s disease and pregnancy complications (3–5).

During the progression from health to periodontal disease, the oral microbiota undergoes a major transition wherein the microbial community increases in total bacterial diversity, accompanied by a bloom in disease-associated bacteria which start to dominate the community, while being otherwise present in low numbers in healthy microbiota (6, 7). *Porphyromonas gingivalis*, a black-pigmenting anaerobic rod bacterium, is well established as one of the organisms which is positively associated with this shift to a dysbiotic microbiota in disease. Indeed, *P. gingivalis* has been shown to act as a trigger for the conversion of the subgingival microbiota to a more diverse and more pathogenic community in animal models of disease (6, 7). This keystone pathogen behaviour has been attributed to the expression of a variety of extracellular virulence determinants including those trafficked through a Type IX secretion system (T9SS) and through the production of large numbers of outer membrane vesicles, which are collectively able to destabilise the host-microbe homeostasis at this mucosal surface (8, 9).

Hemin, iron (III) protoporphyrin IX chloride, is a major source of iron for a variety of organisms, including *P. gingivalis* where it is an essential cofactor for growth. *P. gingivalis* has a dedicated transport system to ensure an efficient accumulation of hemin on the surface for iron utilisation. Subsequent conversion of hemin to μ-oxo bis-haem, and its complexation with iron III, leads to black pigmentation of the bacterial cells, a prominent characteristic of *P. gingivalis* on solid media containing blood (10). *P. gingivalis*, HmuY and HusA are dedicated, surface-exposed hemophore-like proteins that scavenge and bind hemin, free or released from heme containing proteins by gingipain proteases. The complexes deliver the bound heme to outer-membrane receptors for internalization (HmuR or HusB) and heme is then imported through a dedicated TonB-dependent transport machinery (11).

The concentration of environmental hemin acts as a regulator of virulence in this bacterium: cells cultured in high concentrations of hemin - comparable to the situation in a diseased periodontium with elevated bleeding - upregulate the expression of a wide variety of virulence determinants, including a range of hydrolytic enzymes such as the gingipain cysteine proteases, and other protein products many of unknown function (12, 13). High-hemin grown cells are significantly more pathogenic in animal models of tissue destruction compared to cells grown under hemin limitation (8, 14–16). Hence, the availability of hemin in the growth media is critical to the full expression of virulence of *P. gingivalis* and hemin may be regarded as a global regulator of transcription in this bacterium. Although previous studies have characterised the impact of hemin availability on gene expression and virulence in *P. gingivalis*, the regulatory processes underlying these functional changes are not well understood. However, the P*. gingivalis* ATCC33277– genome encodes seven two-component signal transduction system (TCS) pairs of a response regulator and a histidine kinase traditionally involved in gene regulation in other organisms (17). Recently it has been reported that one of these TCS, PorX/PorY, responds to environmental hemin and through the co-ordinated action of PorX and SigP regulates transcription of the T9SS and hemin accumulation; mutants defective in either *porX* or *porY* are attenuated in a mouse model of virulence (18, 19). However, given the magnitude of changes in response to the concentration of hemin, there may be other mechanisms of hemin regulated gene expression in *P. gingivalis*.

Like eukaryotic genomes, bacterial genomes are subject to epigenetic modifications, specifically DNA methylation, which has multiple roles in bacteria including the regulation of transcription. Bacterial DNA methylation includes modifications N6-methyladenine (6mA), 5-methylcytosine (5mC), and N4- methylcytosine (4mC), of which 6mA is most prevalent in prokaryotes. Recent development in single molecule sequencing technologies, for example Nanopore sequencing, allow for the detection of these multiple DNA methylation base modifications. Using this approach, bacterial DNA methylation signatures have recently been shown to impact gene expression profiles and genome stability (20, 21).

In this study, we investigated the impact of hemin availability on the *P. gingivalis* epigenome and transcriptome. *P. gingivalis* W50 was grown in controlled conditions, in continuous culture at a constant pH and identical growth rates, under either limited hemin or excess hemin conditions. Subsequently, DNA sequencing using Nanopore and RNA sequencing using Illumina were carried out, to profile the *P. gingivalis* DNA methylome and transcriptome under each condition. Hemin-dependent differentially modified signatures were identified for gene expression, as well as for 6mA and 5mC DNA methylation profiles both in Dam/Dcm motifs and genome-wide. Comparison of the signals highlighted a cluster of coordinated changes in 6 genes, including in genes related to lactate utilisation and in ABC transporters.

## Results

Gene expression and DNA methylation profiling of *P. gingivalis* W50 was performed after culturing in chemostat runs with excess (5 mg L^-1^) or limited hemin (0.2 mg L^-1^). Biological replicates were collected every 48-hours after achieving steady state, with three biological replicates per experimental condition profiled using Illumina RNA-Seq and Nanopore DNA sequencing technologies (Fig 1).

**Fig 1.**
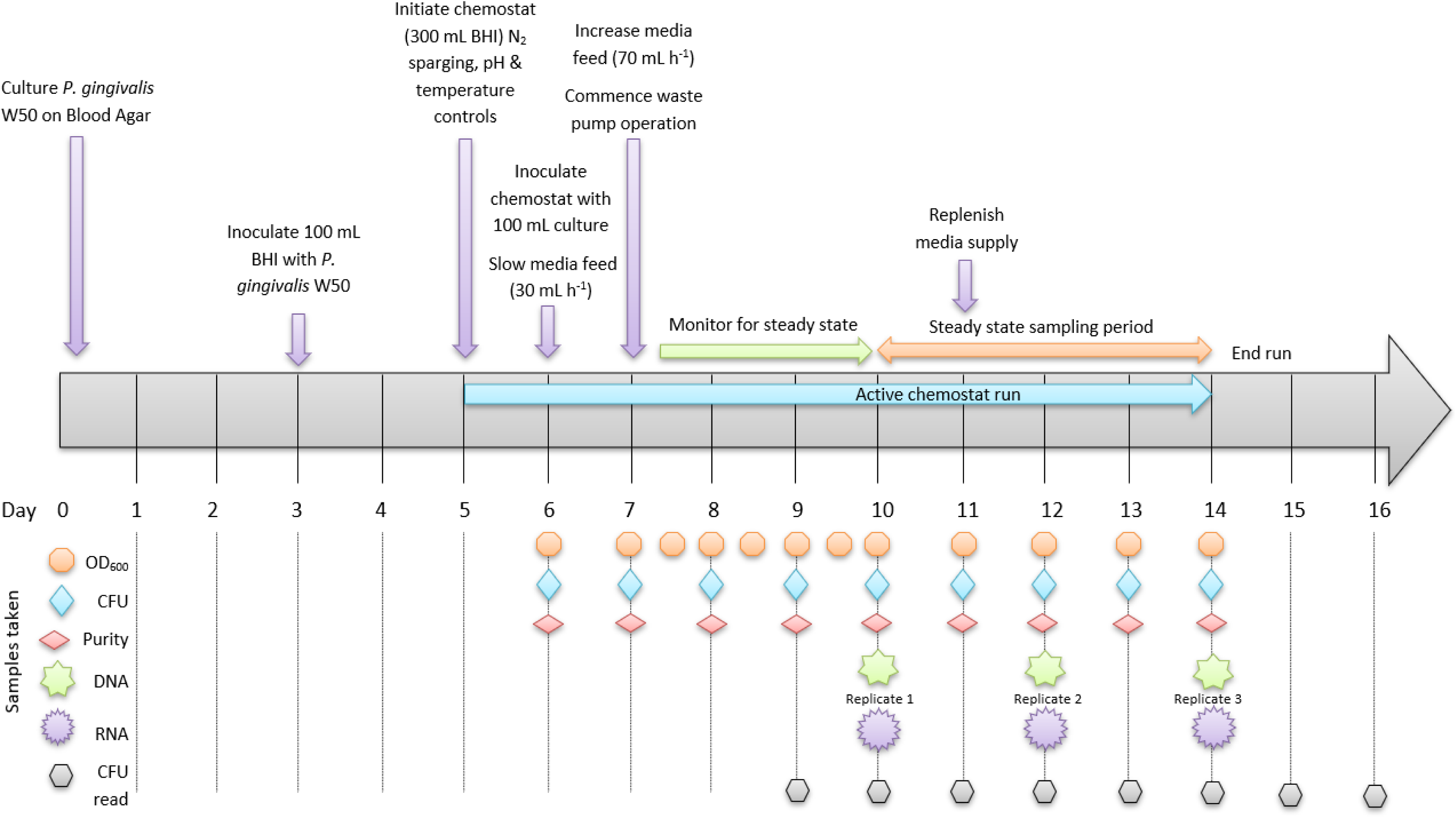
Timeline of *P. gingivalis* W50 chemostat operation and experimental design. CFU – colony forming unit, BHI – Brain Heart Infusion.

### Continuous chemostat environment

Steady state was achieved by days 3 and 4 for excess and limited hemin (Fig 2), respectively. Colony counts of *P. gingivalis* W50 were consistently higher in excess hemin culture versus limitation, elevated by an average of 1.52 × 10^9^ (range 3.44 × 10^8^ – 3.07 × 10^9^) colony forming units per ml (CFU mL^-1^). OD_600_ ranged between 1.67–1.72 (mean 1.69) and 0.357–0.376 (mean 0.367), for excess and limited hemin, respectively. The pH was automatically controlled at pH 7.0 ± 0.1 whilst temperature remained between 36.7 and 37.1°C for both conditions. Bacteria were grown at an identical and constant growth rate of 6.9h mean generation time under both conditions (16).

**Fig 2.**
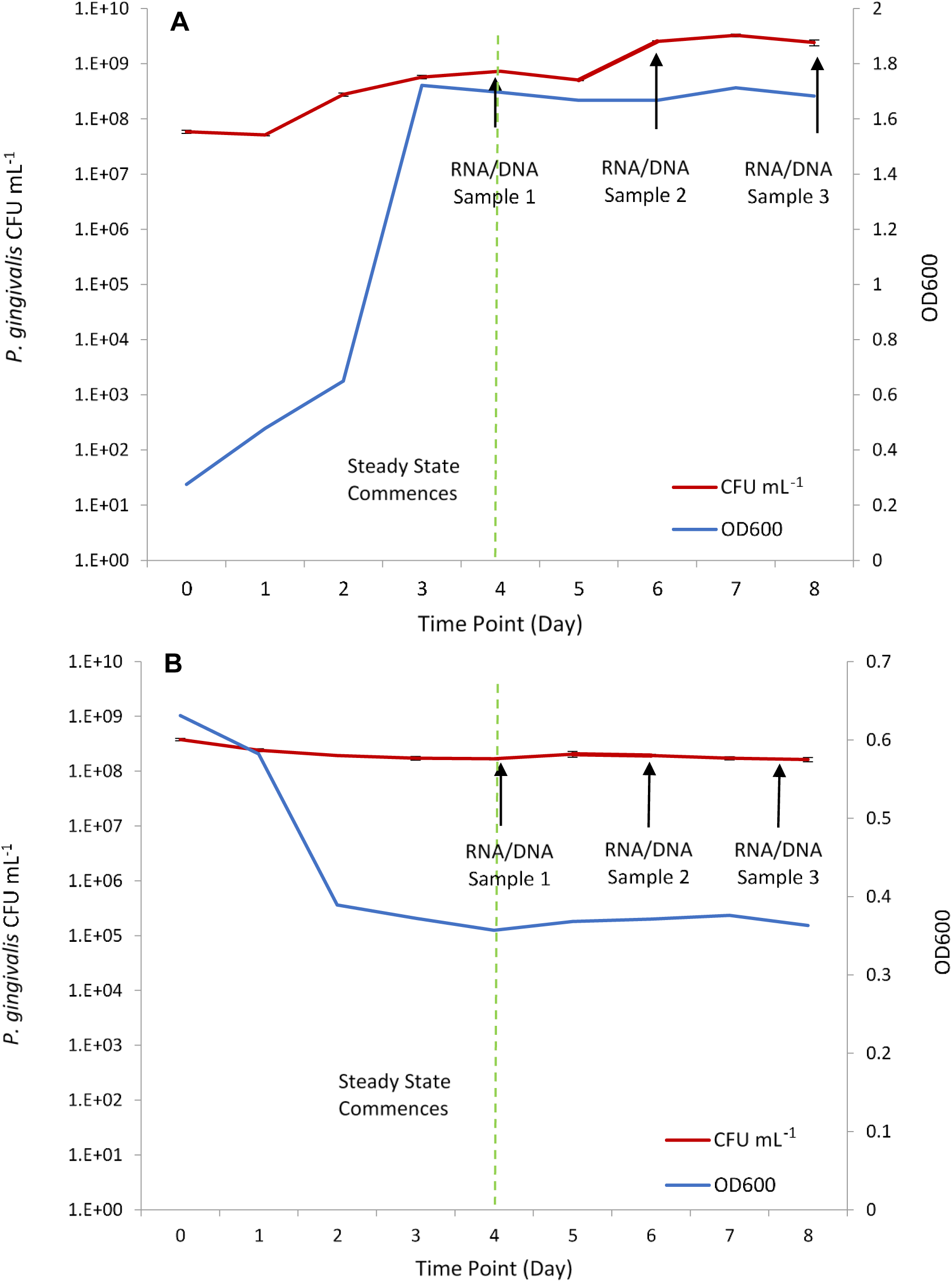
Enumeration of *P. gingivalis* populations throughout the continuous culture chemostat experimental duration by colony count and optical density (OD). A – *P. gingivalis* W50 cultured with excess hemin (5 mg L-1). B – *P. gingivalis* W50 cultured with limited hemin (0.2 mg L-1). CFU – colony forming units. Error bars represent standard error of the mean.

### Gene expression changes

RNA-seq data for the six samples was processed using ProkSeq (22) and DESeq2 (23) to assess gene expression changes observed after culturing *P. gingivalis* W50 in excess (treatment) and limited (control) hemin conditions. These results were mapped to the *P. gingivalis* W50 genome (RefSeq assembly accession no. GCF_000271945.1). Out of 1,192 total genes analysed, those with an expression LFC > 1.5 or LFC < −1.5 were considered differentially expressed after multiple testing correction (FDR = 5%).

Altogether, 161 and 268 differentially expressed genes (DEGs) were respectively over- and under-expressed in excess hemin across the *P. gingivalis* genome (Fig 3). The 161 DEGs over-expressed in excess hemin included genes encoding homologous proteins of the rubrerythrin family (HMPREF1322_RS04040), 4Fe-4S cluster domain-contain containing proteins (HMPREF1322_RS09700, HMPREF1322_RS04860, HMPREF1322_RS04425), a thiamine phosphate synthase (HMPREF1322_RS02110), electron transport complex subunits (HMPREF1322_RS04875, HMPREF1322_RS04865, HMPREF1322_RS01760, HMPREF1322_RS04880, HMPREF1322_RS04870, HMPREF1322_RS04885, HMPREF1322_RS01765), SDR family/ferredoxin oxidoreductases (HMPREF1322_RS07305, HMPREF1322_RS04445, HMPREF1322_RS04440, HMPREF1322_RS00310) and several dehydrogenases (*e.g.* HMPREF1322_RS07315, HMPREF1322_RS04430 and HMPREF1322_RS06505) (Table 1; S1 Table). Methyltransferases targeting tRNA and 16S rRNA molecules were also over-expressed in excess hemin (HMPREF1322_RS00530, HMPREF1322_RS05610, HMPREF1322_RS04605). Most of these genes encode proteins that contain iron as part of their tertiary structure. Three operonic genes, originally labelled as encoding TapA, TapB and TapC (24), HMPREF1322_RS02140, HMPREF1322_RS02145, HMPREF1322_RS02150, respectively, were also over-expressed. TapA, TapB and TapC contribute to virulence in a mouse model as single isogenic mutants are attenuated. Both TapA and TapC exhibit the C-terminal domain signature typical of the T9SS and are therefore cargo proteins as are the proteases HMPREF1322_RS06795 (PrtT) and Lys-gingipain (Kgp).

**Fig 3.**
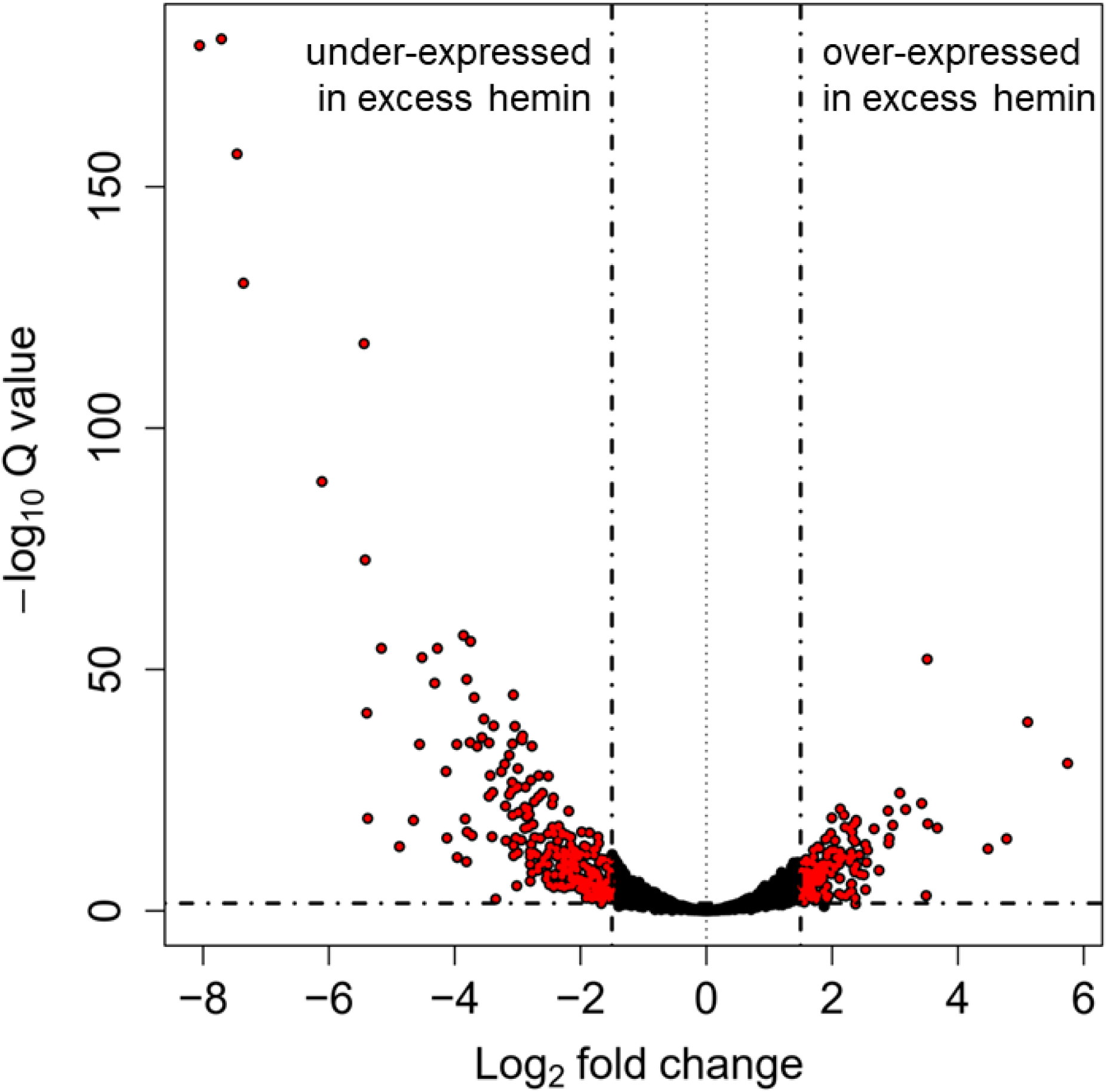
Differentially expressed genes in excess hemin conditions *. P. gingivalis* genes were considered differentially expressed if transcript abundance surpassed 1.5 LFC between culturing conditions (FDR = 5%).

**Table 1.**
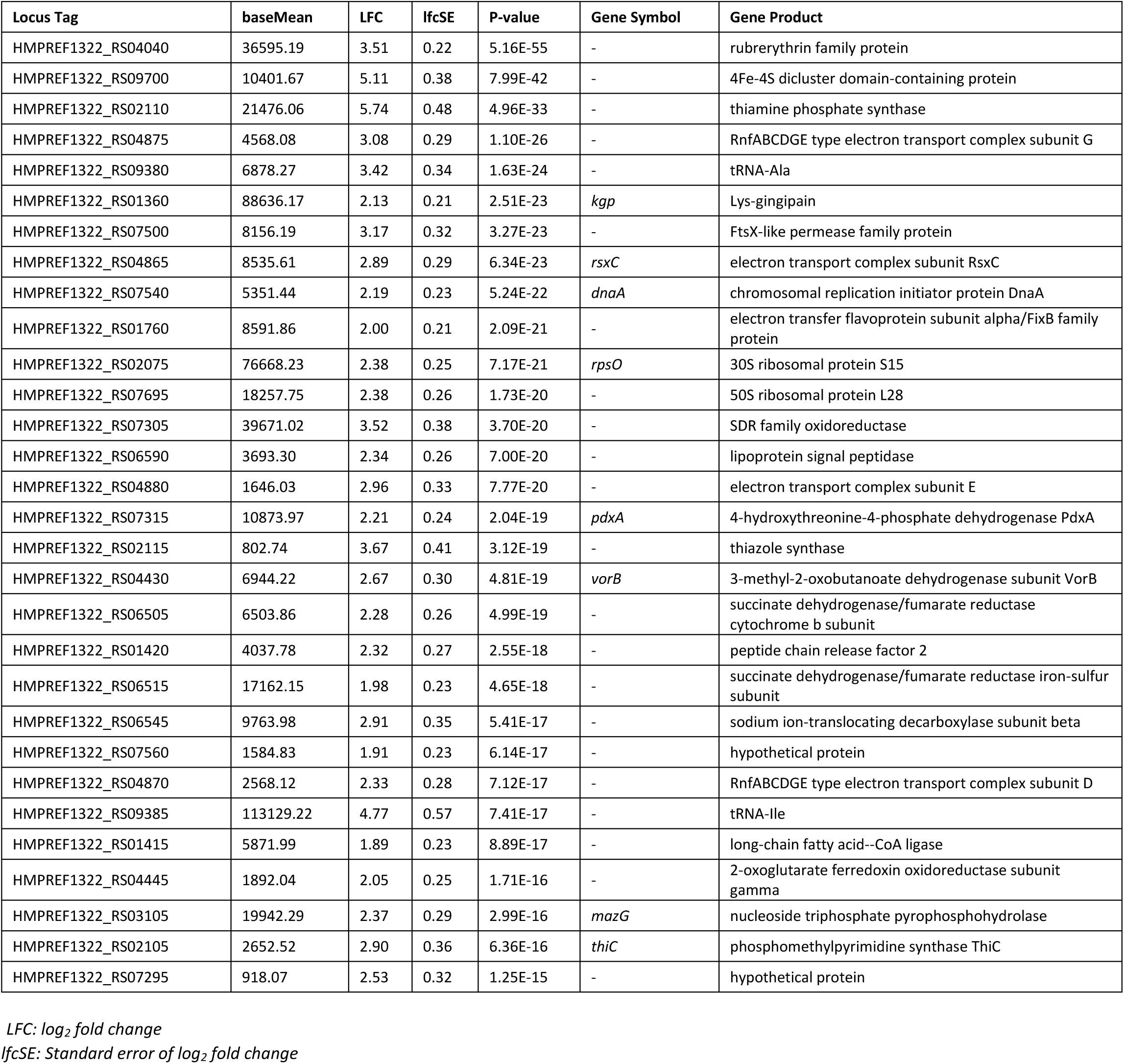
Genes most over-expressed in excess hemin conditions ordered by their significance P -value. The complete list of 161 genes is included in S1 Table.

| Locus Tag | baseMean | LFC | lfcSE | P-value | Gene Symbol | Gene Product |
| --- | --- | --- | --- | --- | --- | --- |
| HMPREF1322_RS04040 | 36595.19 | 3.51 | 0.22 | 5.16E-55 | - | rubrerythrin family protein |
| HMPREF1322_RS09700 | 10401.67 | 5.11 | 0.38 | 7.99E-42 | - | 4Fe-4S dicluster domain-containing protein |
| HMPREF1322_RS02110 | 21476.06 | 5.74 | 0.48 | 4.96E-33 | - | thiamine phosphate synthase |
| HMPREF1322_RS04875 | 4568.08 | 3.08 | 0.29 | 1.10E-26 | - | RnfABCDGE type electron transport complex subunit G |
| HMPREF1322_RS09380 | 6878.27 | 3.42 | 0.34 | 1.63E-24 | - | tRNA-Ala |
| HMPREF1322_RS01360 | 88636.17 | 2.13 | 0.21 | 2.51E-23 | *kgp* | Lys-gingipain |
| HMPREF1322_RS07500 | 8156.19 | 3.17 | 0.32 | 3.27E-23 | - | FtsX-like permease family protein |
| HMPREF1322_RS04865 | 8535.61 | 2.89 | 0.29 | 6.34E-23 | *rsxC* | electron transport complex subunit RsxC |
| HMPREF1322_RS07540 | 5351.44 | 2.19 | 0.23 | 5.24E-22 | *dnaA* | chromosomal replication initiator protein DnaA |
| HMPREF1322_RS01760 | 8591.86 | 2.00 | 0.21 | 2.09E-21 | - | electron transfer flavoprotein subunit alpha/FixB family protein |
| HMPREF1322_RS02075 | 76668.23 | 2.38 | 0.25 | 7.17E-21 | *rpsO* | 30S ribosomal protein S15 |
| HMPREF1322_RS07695 | 18257.75 | 2.38 | 0.26 | 1.73E-20 | - | 50S ribosomal protein L28 |
| HMPREF1322_RS07305 | 39671.02 | 3.52 | 0.38 | 3.70E-20 | - | SDR family oxidoreductase |
| HMPREF1322_RS06590 | 3693.30 | 2.34 | 0.26 | 7.00E-20 | - | lipoprotein signal peptidase |
| HMPREF1322_RS04880 | 1646.03 | 2.96 | 0.33 | 7.77E-20 | - | electron transport complex subunit E |
| HMPREF1322_RS07315 | 10873.97 | 2.21 | 0.24 | 2.04E-19 | *pdxA* | 4-hydroxythreonine-4-phosphate dehydrogenase PdxA |
| HMPREF1322_RS02115 | 802.74 | 3.67 | 0.41 | 3.12E-19 | - | thiazole synthase |
| HMPREF1322_RS04430 | 6944.22 | 2.67 | 0.30 | 4.81E-19 | *vorB* | 3-methyl-2-oxobutanoate dehydrogenase subunit VorB |
| HMPREF1322_RS06505 | 6503.86 | 2.28 | 0.26 | 4.99E-19 | - | succinate dehydrogenase/fumarate reductase cytochrome b subunit |
| HMPREF1322_RS01420 | 4037.78 | 2.32 | 0.27 | 2.55E-18 | - | peptide chain release factor 2 |
| HMPREF1322_RS06515 | 17162.15 | 1.98 | 0.23 | 4.65E-18 | - | succinate dehydrogenase/fumarate reductase iron-sulfur subunit |
| HMPREF1322_RS06545 | 9763.98 | 2.91 | 0.35 | 5.41E-17 | - | sodium ion-translocating decarboxylase subunit beta |
| HMPREF1322_RS07560 | 1584.83 | 1.91 | 0.23 | 6.14E-17 | - | hypothetical protein |
| HMPREF1322_RS04870 | 2568.12 | 2.33 | 0.28 | 7.12E-17 | - | RnfABCDGE type electron transport complex subunit D |
| HMPREF1322_RS09385 | 113129.22 | 4.77 | 0.57 | 7.41E-17 | - | tRNA-Ile |
| HMPREF1322_RS01415 | 5871.99 | 1.89 | 0.23 | 8.89E-17 | - | long-chain fatty acid--CoA ligase |
| HMPREF1322_RS04445 | 1892.04 | 2.05 | 0.25 | 1.71E-16 | - | 2-oxoglutarate ferredoxin oxidoreductase subunit gamma |
| HMPREF1322_RS03105 | 19942.29 | 2.37 | 0.29 | 2.99E-16 | *mazG* | nucleoside triphosphate pyrophosphohydrolase |
| HMPREF1322_RS02105 | 2652.52 | 2.90 | 0.36 | 6.36E-16 | *thiC* | phosphomethylpyrimidine synthase ThiC |
| HMPREF1322_RS07295 | 918.07 | 2.53 | 0.32 | 1.25E-15 | - | hypothetical protein |

*LFC*: log<sub>2</sub> fold change
*lfcSE*: Standard error of log<sub>2</sub> fold change

The 268 DEGs under-expressed in excess hemin included genes *hmuY* (hemin-binding), HMPREF1322_RS06790 (hemin transport), and those encoding homologues of at least 35 transposases (*e.g.* HMPREF1322_RS07370, HMPREF1322_RS07285, HMPREF1322_RS10850) (Table 2; S2 Table). Other genes under-expressed in excess hemin conditions included genes coding several transporter proteins, with at least eight genes coding ABC transporter proteins (*e.g.* HMPREF1322_RS00745, HMPREF1322_RS00115, HMPREF1322_RS08925). Two genes coding for class I SAM-dependent methyltransferases (HMPREF1322_RS07630, HMPREF1322_RS07185) and one gene coding for an isoprenylcysteine carboxylmethyltransferase family protein (HMPREF1322_RS07190) were under-expressed in excess hemin conditions. The gene encoding the largest protein component of the T9SS, Sov (SprA), and also genes encoding OmpH, PorN (GldN) and PorX along with five cargo proteins, are under-expressed in excess hemin (S2 Table).

**Table 2.** Genes most under-expressed in excess hemin conditions (or most over-expressed in limited hemin conditions) ordered by their significance P -value. The complete list of 268 genes is included in S2 Table.

| Locus Tag | baseMean | LFC | lfcSE | P-value | Gene Symbol | Gene Product |
| --- | --- | --- | --- | --- | --- | --- |
| HMPREF1322_RS06000 | 19212.04 | -7.71 | 0.27 | 1.21E-184 | - | TetR/AcrR family transcriptional regulator |
| HMPREF1322_RS06010 | 7387.12 | -8.06 | 0.28 | 5.34E-183 | - | outer membrane lipoprotein-sorting protein |
| HMPREF1322_RS06015 | 24454.31 | -7.46 | 0.28 | 2.46E-160 | - | hypothetical protein |
| HMPREF1322_RS06005 | 47425.12 | -7.36 | 0.30 | 1.87E-133 | - | MMPL family transporter |
| HMPREF1322_RS05430 | 7754.25 | -5.44 | 0.23 | 7.95E-121 | - | flavodoxin |
| HMPREF1322_RS06790 | 15141.08 | -6.11 | 0.30 | 3.67E-92 | - | heme-binding protein HmuY |
| HMPREF1322_RS04145 | 4231.05 | -5.42 | 0.29 | 7.09E-76 | - | DUF4876 domain-containing protein |
| HMPREF1322_RS00165 | 33997.53 | -3.86 | 0.24 | 3.60E-60 | - | hypothetical protein |
| HMPREF1322_RS08955 | 18702.05 | -3.75 | 0.23 | 6.92E-59 | - | PglZ domain-containing protein |
| HMPREF1322_RS03170 | 5510.46 | -5.17 | 0.32 | 2.13E-57 | - | T9SS type A sorting domain-containing protein |
| HMPREF1322_RS00745 | 3619.54 | -4.27 | 0.27 | 2.40E-57 | - | ABC transporter ATP-binding protein |
| HMPREF1322_RS00465 | 1031.56 | -4.52 | 0.29 | 2.04E-55 | - | DUF3575 domain-containing protein |
| HMPREF1322_RS03855 | 47852.31 | -3.81 | 0.25 | 8.15E-51 | - | T9SS type A sorting domain-containing protein |
| HMPREF1322_RS05045 | 1138.34 | -4.32 | 0.29 | 5.25E-50 | - | hypothetical protein |
| HMPREF1322_RS07370 | 25172.29 | -3.07 | 0.21 | 1.50E-47 | - | transposase |
| HMPREF1322_RS05050 | 901.14 | -3.69 | 0.26 | 5.74E-47 | - | hypothetical protein |
| HMPREF1322_RS04140 | 4263.72 | -5.40 | 0.39 | 9.54E-44 | - | TonB-dependent receptor |
| HMPREF1322_RS07285 | 4362.71 | -3.54 | 0.26 | 1.81E-42 | - | IS5/IS1182 family transposase |
| HMPREF1322_RS00730 | 1091.42 | -3.38 | 0.25 | 5.01E-41 | - | hypothetical protein |
| HMPREF1322_RS10850 | 767.65 | -3.04 | 0.23 | 6.51E-41 | - | IS5/IS1182 family transposase |
| HMPREF1322_RS05700 | 1363.75 | -2.92 | 0.22 | 6.42E-39 | - | IS5/IS1182 family transposase |
| HMPREF1322_RS02830 | 245.42 | -3.57 | 0.27 | 1.53E-38 | - | helix-turn-helix domain-containing protein |
| HMPREF1322_RS08255 | 1559.13 | -2.94 | 0.23 | 2.65E-38 | - | IS5/IS1182 family transposase |
| HMPREF1322_RS00725 | 911.64 | -2.93 | 0.23 | 5.26E-38 | - | lactate utilization protein |
| HMPREF1322_RS05425 | 530.83 | -3.76 | 0.29 | 1.83E-37 | - | DUF2023 family protein |
| HMPREF1322_RS00485 | 425.30 | -3.46 | 0.27 | 2.24E-37 | - | DUF4906 domain-containing protein |
| HMPREF1322_RS07380 | 45140.22 | -3.08 | 0.24 | 3.93E-37 | - | family outer membrane protein |
| HMPREF1322_RS08605 | 2232.06 | -4.56 | 0.36 | 4.87E-37 | - | co-chaperone GroES |
| HMPREF1322_RS07630 | 1708.56 | -3.97 | 0.31 | 5.27E-37 | - | class I SAM-dependent methyltransferase |
| HMPREF1322_RS04045 | 1062.32 | -2.77 | 0.22 | 1.27E-36 | - | IS982 family transposase |

*LFC: log<sub>2</sub> fold change*
*lfcSE: Standard error of log<sub>2</sub> fold change*

Different types of genes involved in virulence were identified in both the under- and over-expressed DEG subsets. For example, these included different types of N-acetylmuramoyl-L-alanine amidase genes, T9SS type A sorting domain-containing protein genes and MATE family efflux transporter genes, and tetratricopeptide repeat protein genes were identified (S1 and S2 Tables).

We explored molecular functions shared across the DEGs, although a proportion of genes were not well characterised. Sixty-seven under-expressed (25% of total) and 19 over-expressed (12%) genes identified during growth in excess hemin encoded hypothetical proteins. Additionally, only 16 (6%) of under-expressed genes and 49 (30%) of over-expressed genes identified had well-characterised annotations in RefSeq. A GO term enrichment analysis was carried out using ShinyGO and gene symbols that matched genes in the reference strain *P. gingivalis* W83 (25), with altogether 8 (out of 16 well-characterised) under-expressed and 40 (out of 49 well-characterised) over-expressed genes.

Forty-two and 43 GO molecular functions were respectively enriched in genes under- and over-expressed in excess hemin conditions. In both cases, molecular functions identified were mostly related to binding activities such as cyclic compound binding, heterocyclic compound binding and ion binding (Fig 4; S3 and S4 Tables). In excess hemin, functions related to Fe-S cluster binding, 4Fe-4S cluster binding and metal cluster binding, and several different catalytic activities, including GTPase, metallo-endopeptidase, active transmembrane transporter and diphosphotransferase activities were also identified (Fig 4; S4 Table).

**Fig 4.**
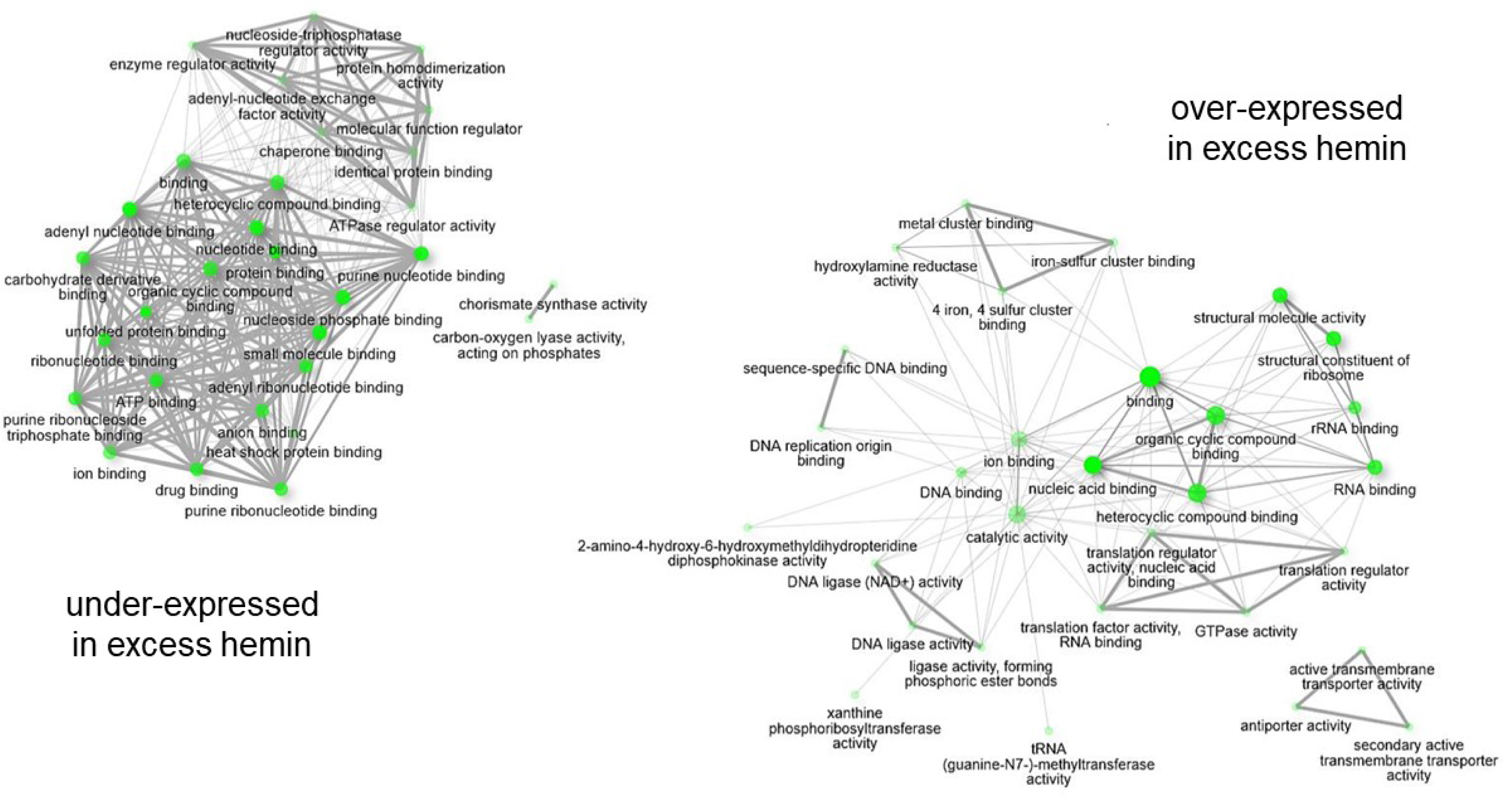
GO molecular function enrichment of genes under an d over-expressed in excess hemin conditions (1.5 LFC, FDR = 10%). The top 30 GO molecular functions were selected to create the networks and edges represent pathways with > 10% shared genes. Darker nodes are more significantly enriched, bigger nodes repres ent larger gene sets and thicker edges correspond to more overlap s between gene sets. Pathways were created based on gene matches to *P. gingivalis* W83 reference genome.

Two sensitivity analyses were carried out for the genome-wide DESeq2 DEG results presented here. A first sensitivity analysis was performed using edgeR, which normalizes expression data differently to DESeq2, and consistently identified the majority of DEGs identified with DESeq2 (S1 Fig). Specifically, edgeR identified 166 genes as over-expressed and 266 genes as under-expressed with excess hemin, of which 161 and 265 were previously identified with DESeq2. In a second general sensitivity analysis, a principal component analysis (PCA) of the genome-wide RNA-seq levels also confirmed overall global differences in gene expression levels between the culturing conditions (S2 Fig).

### DNA methylation changes

Nanopore sequencing data were generated for the same six biological samples analysed with Illumina RNA sequencing. Dam and Dcm-dependent methylation was characterised using the Guppy basecaller. A total of 12,066 ‘GATC’ (Dam) and 819 ‘CCWGG’ (Dcm) motifs with minimum 10× coverage across samples were considered. Motif-independent methylation levels for 6mA and 5mC were characterised using Tombo with ‘all-context’ models, considering >2M positions with a minimum 10× coverage. PCA of the genome-wide DNA methylation levels pointed to both global 6mA and 5mC differences in *P. gingivalis* according to hemin culture conditions (S3-A Fig) and mirrored the PCA pattern observed for gene expression (S1 Fig). Similar results were observed when considering signals with at least 100× coverage (S3-B Fig), and Dam-dependent adenine methylation alone (S4-A and S4-B Figs, top panels). Differential methylation analyses with respect to hemin levels were then carried out, considering signals where at least 5% mean methylation difference between the experimental conditions was observed after multiple testing correction (FDR 5%).

### Dam/Dcm DNA methylation

DNA methylation signals were analysed at 79 ‘GATC’ motifs in the genome of *P. gingivalis* W50, which had at least 5% mean methylation differences between hemin conditions. Of these, 36 motifs showed statistically significant differential methylation signals across hemin conditions (S5 Table). Differentially methylated ‘GATC’ motifs (‘GATC’-DMMs) were located in altogether 32 genes, including a zinc-dependent metalloprotease (HMPREF1322_RS05790), a thioredoxin-disulfide reductase (HMPREF1322_RS01475) and a 4-alpha-glucanotransferase (HMPREF1322_RS03650). If genes up to ±1 kb away from the ‘GATC’ motifs were considered, 85 genes were candidates for possible Dam-like regulation. Sensitivity analysis at 100× coverage (66 motifs) identified 36 differentially methylated motifs as well, of which 34 were present in the main analysis (S6 Table).

No minimum 5% mean methylation difference was observed at each Dcm-associated ‘CCWGG’ motif, in line with previous observation for no overall differences in the Dcm-based PCA analysis (S4-A and S4-B Figs, bottom panels). This suggests either the absence of enzymes targeting ‘CCWGG’ motifs in *P. gingivalis* W50 or a minimal role of a Dcm-like response to hemin availability.

### All-context 6mA/5mC DNA methylation

The proportion of differential methylation at each genomic position was analysed for altogether 139,794 adenines and 87,398 cytosines with at least 5% mean difference in DNA methylation levels between conditions.

Forty-nine differentially methylated adenine sites (DMAs) were identified between experimental conditions across the *P. gingivalis* W50 genome (Fig 5). A cluster of 15 DMAs was observed in a genomic region (NZ_AJZS01000011.1 ≈ 12 – 20 kb) containing the (Fe-S)-binding protein HMPREF1322_RS00720, lactate utilization protein HMPREF1322_RS00725, hypothetical protein HMPREF1322_RS00730, PaaI family thioesterase HMPREF1322_RS00735 and ABC transporter ATP-binding protein HMPREF1322_RS00740 (Fig 5 top panel, purple; Fig 6). In annotating DMAs to genes, if genes up to ±1 kb away from the DMA on both strands were considered, all DMAs are annotated to at least one gene for a total of 75 genes altogether. Furthermore, 25 DMAs were annotated to genes on the same strand, including seven DMAs encoding a lactate utilization protein (HMPREF1322_RS00725), five in a hypothetical protein (HMPREF1322_RS00730), and two each in a 4-alpha-glucanotransferase (HMPREF1322_RS03650) and a Ppx/GppA family phosphatase (HMPREF1322_RS06180) (Table 3). Four DMAs were located within 1 kb of a gene encoding a (Fe-S)-binding protein (HMPREF1322_RS00720). In a sensitivity analysis considering only adenines with at least 100× coverage (85,090 adenines), 20 DMAs were identified at FDR = 5% (S7 Table) and all 20 were also identified in the results of the main analysis.

**Fig 5.**
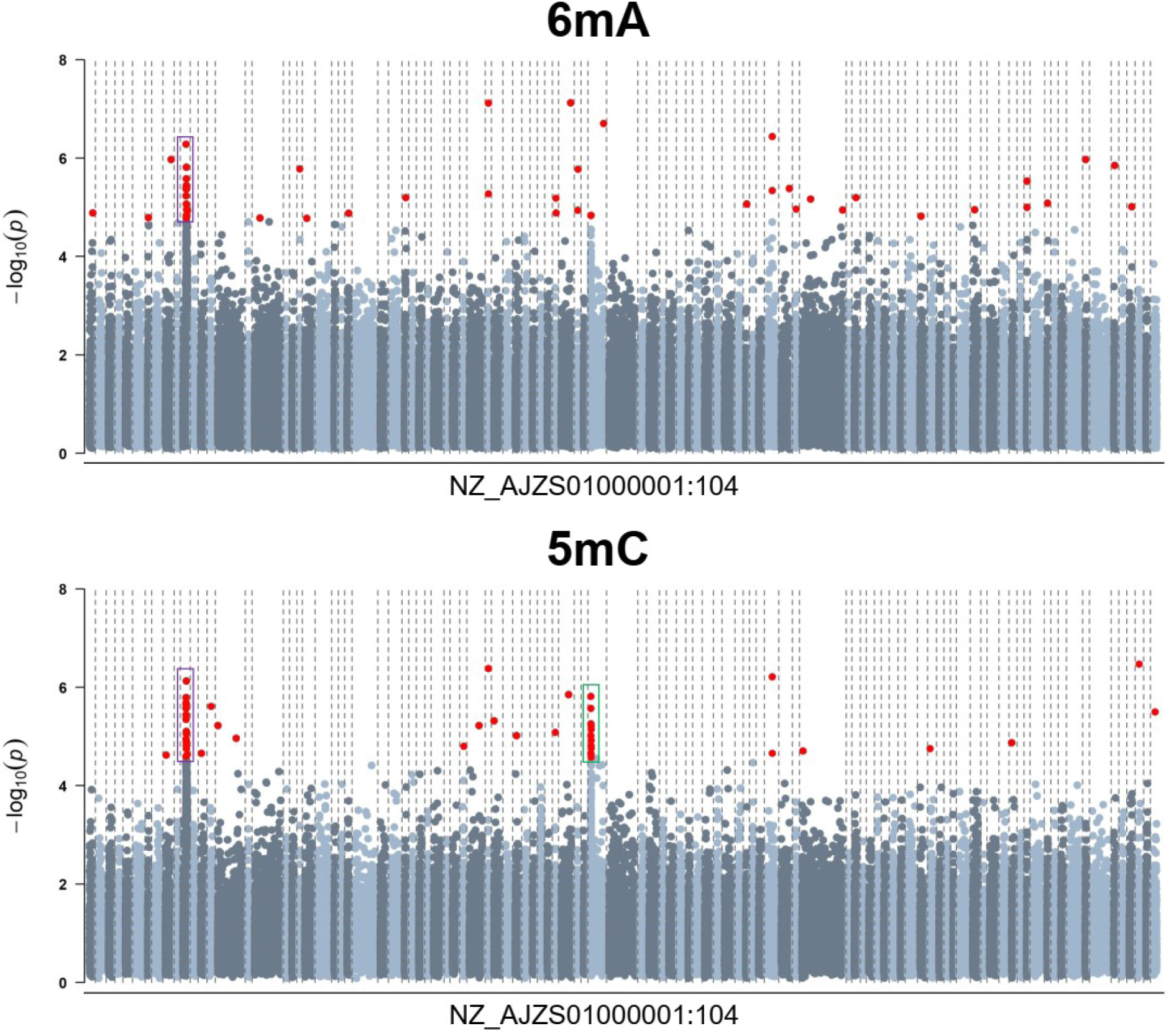
Differential methylation analysis results for excess hemin. Results are presented for 139,794 adenines (6mA) and 87,398 cytosines (5mC) showing a minimum 5% mean methylation difference between experimental conditions. Contigs were plotted in ascending order (NZ_AJZS01000001:104) and are delimited by the dashed lines in the graph. Genomic positions surpassing the FDR = 5% significance threshold are depicted in red. Clusters of DMAs and DMCs are highlighted in purple and green. The DMA/DMC cluster highlighted in purple is shown in detail in Fig 6.

**Fig 6.**
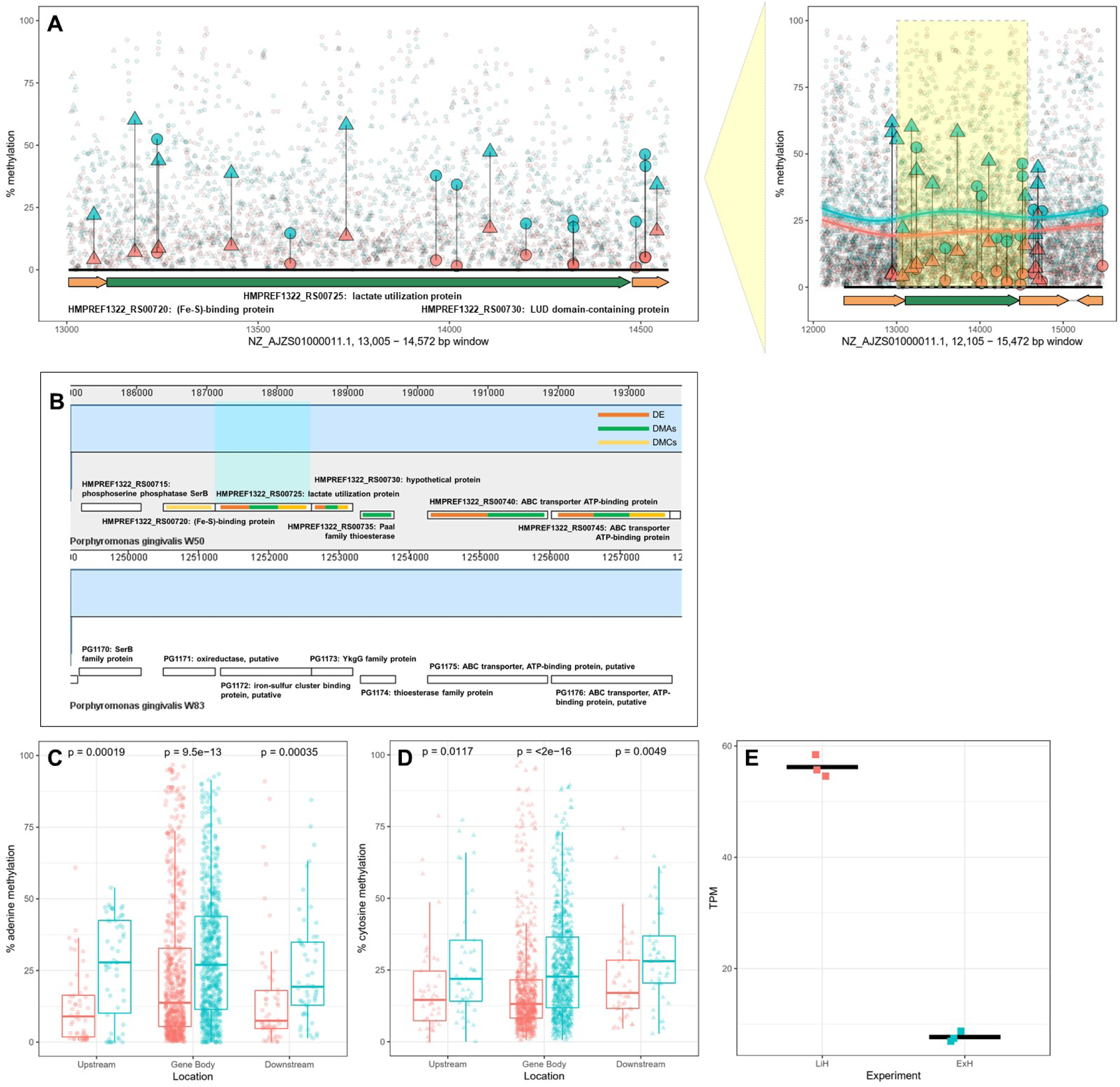
Genomic cluster of differential expression, adenine and cytosine methylation in the lactate utilization protein (HMPREF1322_RS00725) gene locus. Adenine (circles) and cytosine (triangles) percentage of methylation in this gene locus are shown in limited (LiH; red) and excess (ExH; blue) hemin conditions. DMAs and DMCs (larger data points) were observed within a 100 bp or 1000 bp window from the gene and a generalized additive model curve was fitted to the methylation values (A). The lactate utilization protein HMPREF1322_RS00725 was located in a cluster of genes conserved between *P. gingivalis* W50 and reference strain W83 (B). Genes in this location were affected by both DNA methylation (DMA/DMC) and gene expression (DE) changes. Differential adenine (C) and cytosine (D) methylation was statistically significant upstream (−100 bp), downstream (+100 bp) and in the gene body of HMPREF1322_RS00725 using the Wilcoxon signed-ranked test. Increased methylation in ExH was accompanied by decreased gene expression of HMPREF1322_RS00725 (E).

**Table 3.**
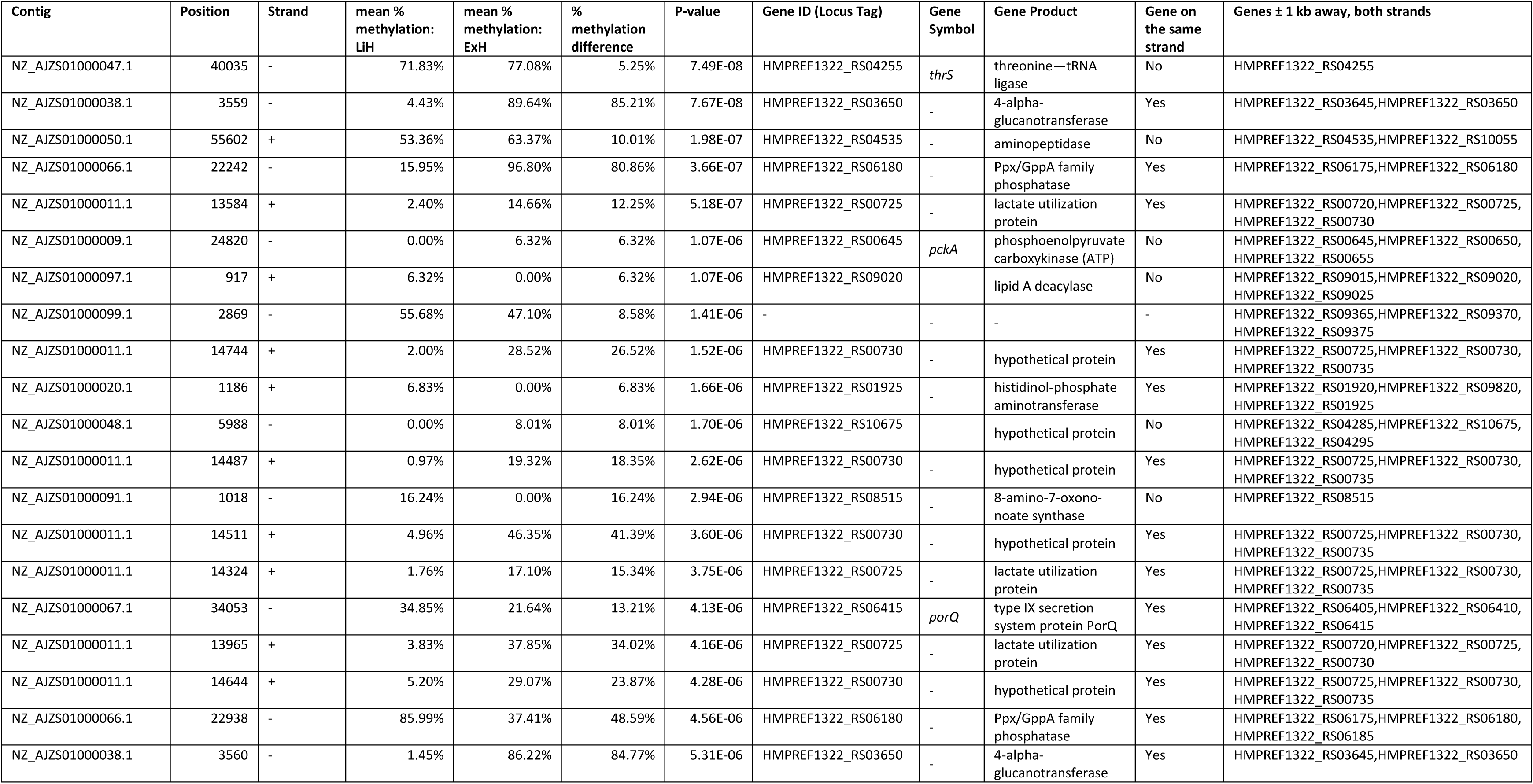
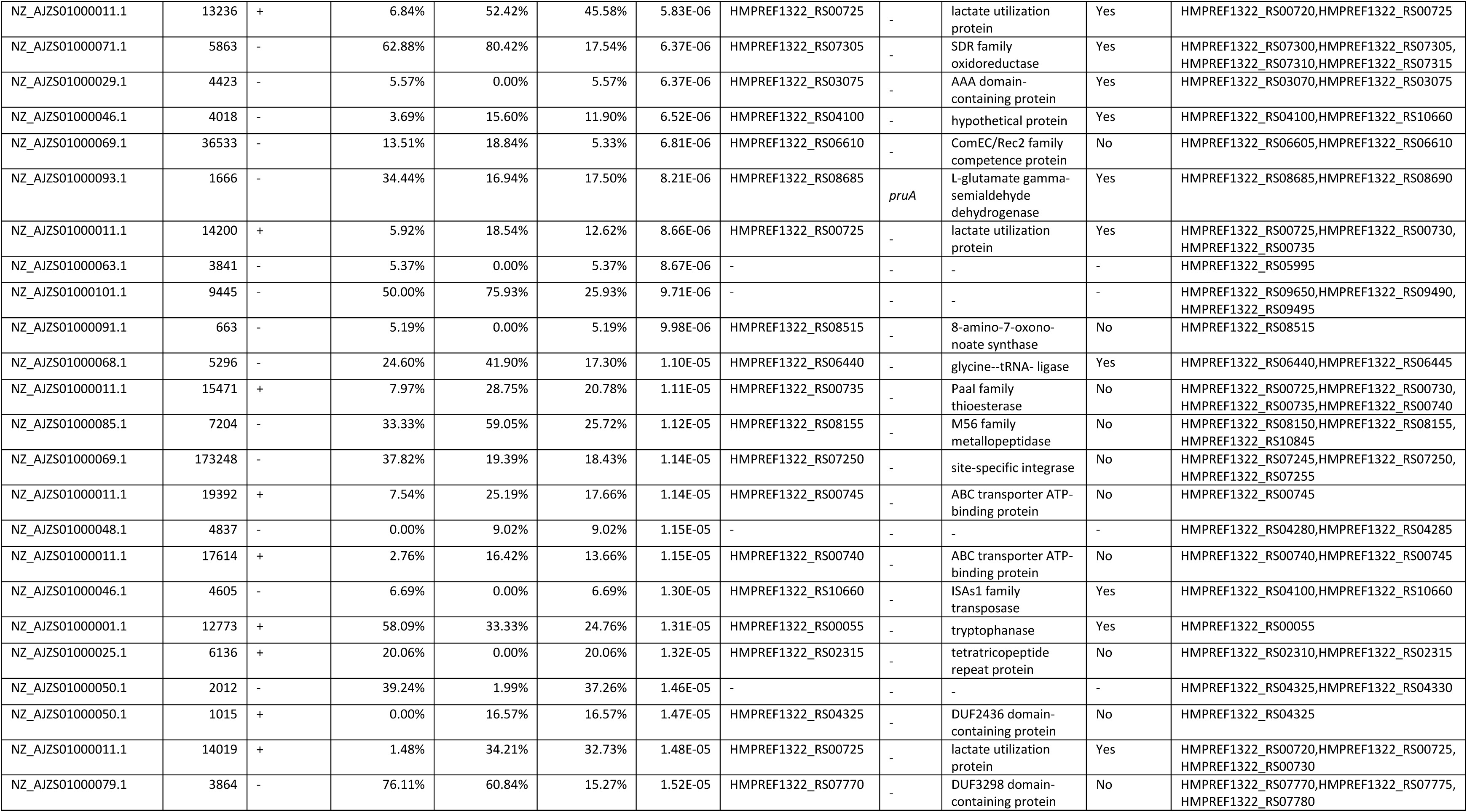
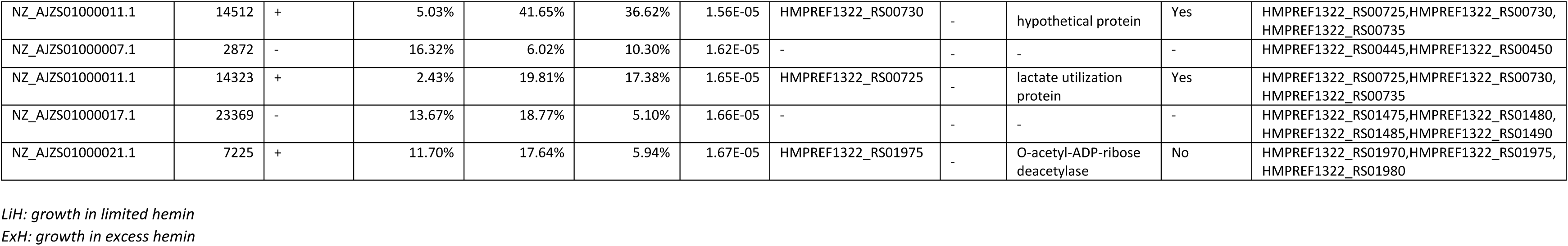
DMAs (6mA) identified in *P. gingivalis* W50, selecting for a minimum mean 5% methylation difference between experimental conditions (FDR = 5% and minimum 10× coverage).

Forty-seven differentially methylated cytosines (DMCs) were identified between experimental conditions across the *P. gingivalis* W50 genome (Fig 5). Two clusters of 15 and 12 DMCs emerged - the first was the same as the DMA cluster region (NZ_AJZS01000011.1, ≈ 12 – 20 kb), and the second was in a genomic region (NZ_AJZS01000050.1, ≈ 0 – 3.5 kb) containing the DUF2436 domain-containing protein HMPREF1322_RS04325 and nucleoside permease HMPREF1322_RS04330 (Fig. 5 bottom panel, purple and green; Fig 6). Thirty-five DMCs were located in 16 genes including in genes identified in the DMA analyses above. Specifically, four DMCs were located in genes encoding the (Fe-S)-binding protein HMPREF1322_RS00720, two in the lactate utilization protein HMPREF1322_RS00725, and two in the hypothetical protein HMPREF1322_RS00730 reported above for 6mA. Single DMCs were also identified in the 4-alpha-glucanotransferase HMPREF1322_RS03650 and Ppx/GppA family phosphatase HMPREF1322_RS06180 genes (Table 4). Overall, if genes up to ±1 kb away on both strands were considered, 31 genes harboured DMCs, of which 10 also harboured DMAs. Other genes annotated to the DMCs identified included those encoding transporter proteins such as an iron ABC transporter permease (HMPREF1322_RS01240) and others (HMPREF1322_RS01245; HMPREF1322_RS06460). In sensitivity analysis considering only cytosines with at least 100× coverage (46,930 cytosines), 47 DMCs were also identified, of which 27 were also in the main analysis results (S8 Table).

**Table 4.** DMCs (5mC) identified in *P. gingivalis* W50, selecting for a minimum mean 5% methylation difference between experimental conditions (FDR = 5% and minimum 10× coverage).

| Contig | Position | Strand | mean % methylation: LiH | mean % methylation: ExH | % methylation difference | P-value | Gene ID (Locus Tag) | Gene Symbol | Gene Product | Gene on the same strand | Genes ± 1 kb away, both strands |
| --- | --- | --- | --- | --- | --- | --- | --- | --- | --- | --- | --- |
| NZ_AJZS01000102.1 | 6840 | - | 6.63% | 12.27% | 5.64% | 3.39E-07 | HMPREF1322_RS09535 | - | pyridoxal phosphate-dependent aminotransferase | Yes | HMPREF1322_RS09535,HMPREF1322_RS09540 |
| NZ_AJZS01000038.1 | 3558 | - | 5.53% | 95.22% | 89.69% | 4.16E-07 | HMPREF1322_RS03650 | - | 4-alpha-glucanotransferase | Yes | HMPREF1322_RS03645,HMPREF1322_RS03650 |
| NZ_AJZS01000066.1 | 22243 | - | 10.21% | 64.13% | 53.91% | 6.11E-07 | HMPREF1322_RS06180 | - | Ppx/GppA family phosphatase | Yes | HMPREF1322_RS06175,HMPREF1322_RS06180 |
| NZ_AJZS01000011.1 | 14106 | + | 16.77% | 47.36% | 30.59% | 7.45E-07 | HMPREF1322_RS00725 | - | lactate utilization protein | Yes | HMPREF1322_RS00720,HMPREF1322_RS00725,HMPREF1322_RS00730 |
| NZ_AJZS01000047.1 | 30511 | + | 65.91% | 58.49% | 7.42% | 1.42E-06 | HMPREF1322_RS04210 | - | S9 family peptidase | Yes | HMPREF1322_RS04205,HMPREF1322_RS04210 |
| NZ_AJZS01000050.1 | 1612 | - | 34.96% | 8.22% | 26.74% | 1.53E-06 | HMPREF1322_RS04325 | - | DUF2436 domain-containing protein | Yes | HMPREF1322_RS04325,HMPREF1322_RS04330 |
| NZ_AJZS01000011.1 | 13240 | + | 8.76% | 43.87% | 35.12% | 1.61E-06 | HMPREF1322_RS00725 | - | lactate utilization protein | Yes | HMPREF1322_RS00720,HMPREF1322_RS00725 |
| NZ_AJZS01000011.1 | 12946 | + | 4.71% | 58.08% | 53.37% | 2.09E-06 | HMPREF1322_RS00720 | - | (Fe-S)-binding protein | Yes | HMPREF1322_RS00715,HMPREF1322_RS00720,HMPREF1322_RS00725 |
| NZ_AJZS01000011.1 | 13178 | + | 7.08% | 60.16% | 53.08% | 2.24E-06 | HMPREF1322_RS00725 | - | lactate utilization protein | Yes | HMPREF1322_RS00720,HMPREF1322_RS00725 |
| NZ_AJZS01000014.1 | 5938 | + | 34.76% | 52.63% | 17.87% | 2.45E-06 | HMPREF1322_RS00860 | htpG | molecular chaperone HtpG | No | HMPREF1322_RS00860 |
| NZ_AJZS01000050.1 | 1569 | - | 53.78% | 2.75% | 51.03% | 2.69E-06 | HMPREF1322_RS04325 | - | DUF2436 domain-containing protein | Yes | HMPREF1322_RS04325,HMPREF1322_RS04330 |
| NZ_AJZS01000011.1 | 12945 | + | 5.18% | 61.79% | 56.61% | 2.71E-06 | HMPREF1322_RS00720 | - | (Fe-S)-binding protein | Yes | HMPREF1322_RS00715,HMPREF1322_RS00720,HMPREF1322_RS00725 |
| NZ_AJZS01000104.1 | 8834 | - | 0.00% | 7.90% | 7.90% | 3.17E-06 | HMPREF1322_RS09610 | - | tetratricopeptide repeat protein | Yes | HMPREF1322_RS09605,HMPREF1322_RS09610 |
| NZ_AJZS01000011.1 | 13002 | + | 3.76% | 55.38% | 51.62% | 3.63E-06 | HMPREF1322_RS00720 | - | (Fe-S)-binding protein | Yes | HMPREF1322_RS00715,HMPREF1322_RS00720,HMPREF1322_RS00725 |
| NZ_AJZS01000011.1 | 13071 | + | 4.20% | 21.99% | 17.79% | 4.48E-06 | HMPREF1322_RS00720 | - | (Fe-S)-binding protein | Yes | HMPREF1322_RS00720,HMPREF1322_RS00725 |
| NZ_AJZS01000039.1 | 305 | + | 6.19% | 17.19% | 10.99% | 4.82E-06 | - | - | - | - | HMPREF1322_RS03655 |
| NZ_AJZS01000050.1 | 1624 | - | 31.26% | 11.13% | 20.13% | 5.57E-06 | HMPREF1322_RS04325 | - | DUF2436 domain-containing protein | Yes | HMPREF1322_RS04325,HMPREF1322_RS04330 |
| NZ_AJZS01000050.1 | 2099 | - | 58.19% | 8.09% | 50.10% | 6.00E-06 | HMPREF1322_RS04330 | - | nucleoside permease | Yes | HMPREF1322_RS04325,HMPREF1322_RS04330 |
| NZ_AJZS01000037.1 | 41404 | - | 62.40% | 68.48% | 6.08% | 6.02E-06 | HMPREF1322_RS03585 | - | CusA/CzcA family heavy metal efflux RND transporter | No | HMPREF1322_RS03585,HMPREF1322_RS03590 |
| NZ_AJZS01000015.1 | 952 | + | 13.70% | 22.02% | 8.33% | 6.03E-06 | HMPREF1322_RS00885 | - | U32 family peptidase | No | HMPREF1322_RS00880,HMPREF1322_RS00885 |
| NZ_AJZS01000050.1 | 1920 | - | 64.72% | 8.39% | 56.32% | 7.23E-06 | - | - | - | - | HMPREF1322_RS04325,HMPREF1322_RS04330 |
| NZ_AJZS01000011.1 | 14698 | + | 26.82% | 44.78% | 17.96% | 7.90E-06 | HMPREF1322_RS00730 | - | hypothetical protein | Yes | HMPREF1322_RS00725,HMPREF1322_RS00730,<br>HMPREF1322_RS00735 |
| NZ_AJZS01000046.1 | 1044 | - | 10.78% | 5.27% | 5.52% | 8.29E-06 | HMPREF1322_RS04090 | - | MBOAT family protein | Yes | HMPREF1322_RS04090,HMPREF1322_RS04095 |
| NZ_AJZS01000011.1 | 14542 | + | 15.67% | 34.20% | 18.53% | 8.76E-06 | HMPREF1322_RS00730 | - | hypothetical protein | Yes | HMPREF1322_RS00725,HMPREF1322_RS00730,<br>HMPREF1322_RS00735 |
| NZ_AJZS01000041.1 | 4471 | - | 17.52% | 22.93% | 5.41% | 9.62E-06 | - | - | - | - | HMPREF1322_RS03860,HMPREF1322_RS03870,<br>HMPREF1322_RS03875 |
| NZ_AJZS01000050.1 | 1625 | - | 26.00% | 9.52% | 16.48% | 9.79E-06 | HMPREF1322_RS04325 | - | DUF2436 domain-containing protein | Yes | HMPREF1322_RS04325,HMPREF1322_RS04330 |
| NZ_AJZS01000015.1 | 78713 | - | 64.82% | 58.15% | 6.67% | 1.08E-05 | HMPREF1322_RS01240 | - | iron ABC transporter<br>permease | Yes | HMPREF1322_RS01240,HMPREF1322_RS01245 |
| NZ_AJZS01000011.1 | 13730 | + | 13.59% | 58.15% | 44.56% | 1.13E-05 | HMPREF1322_RS00725 | - | lactate utilization protein | Yes | HMPREF1322_RS00720,HMPREF1322_RS00725,<br>HMPREF1322_RS00730 |
| NZ_AJZS01000050.1 | 2262 | - | 53.40% | 9.81% | 43.59% | 1.20E-05 | HMPREF1322_RS04330 | - | nucleoside permease | Yes | HMPREF1322_RS04325,HMPREF1322_RS04330 |
| NZ_AJZS01000011.1 | 14669 | + | 7.23% | 19.84% | 12.61% | 1.28E-05 | HMPREF1322_RS00730 | - | hypothetical protein | Yes | HMPREF1322_RS00725,HMPREF1322_RS00730,<br>HMPREF1322_RS00735 |
| NZ_AJZS01000089.1 | 26 | + | 0.00% | 9.03% | 9.03% | 1.34E-05 | HMPREF1322_RS08455 | - | IS5/IS1182 family<br>transposase | Yes | HMPREF1322_RS08455,HMPREF1322_RS08460 |
| NZ_AJZS01000011.1 | 14706 | + | 2.91% | 21.70% | 18.79% | 1.52E-05 | HMPREF1322_RS00730 | - | hypothetical protein | Yes | HMPREF1322_RS00725,HMPREF1322_RS00730,<br>HMPREF1322_RS00735 |
| NZ_AJZS01000050.1 | 2002 | - | 58.99% | 4.81% | 54.18% | 1.57E-05 | - | - | - | - | HMPREF1322_RS04325,HMPREF1322_RS04330 |
| NZ_AJZS01000036.1 | 5418 | + | 22.76% | 14.46% | 8.30% | 1.59E-05 | HMPREF1322_RS03420 | - | YjgP/YjgQ family<br>permease | No | HMPREF1322_RS03410,HMPREF1322_RS03415,<br>HMPREF1322_RS03420,HMPREF1322_RS03425 |
| NZ_AJZS01000050.1 | 2101 | - | 53.34% | 4.65% | 48.69% | 1.74E-05 | HMPREF1322_RS04330 | - | nucleoside permease | Yes | HMPREF1322_RS04325,HMPREF1322_RS04330 |
| NZ_AJZS01000080.1 | 123 | + | 12.46% | 0.00% | 12.46% | 1.76E-05 | HMPREF1322_RS07835 | - | hypothetical protein | Yes | HMPREF1322_RS07835,HMPREF1322_RS07840,<br>HMPREF1322_RS07845 |
| NZ_AJZS01000080.1 | 124 | + | 12.46% | 0.00% | 12.46% | 1.76E-05 | HMPREF1322_RS07835 | - | hypothetical protein | Yes | HMPREF1322_RS07835,HMPREF1322_RS07840,<br>HMPREF1322_RS07845 |
| NZ_AJZS01000011.1 | 14693 | + | 14.07% | 38.73% | 24.67% | 1.77E-05 | HMPREF1322_RS00730 | - | hypothetical protein | Yes | HMPREF1322_RS00725,HMPREF1322_RS00730,<br>HMPREF1322_RS00735 |
| NZ_AJZS01000069.1 | 3434 | - | 51.80% | 41.48% | 10.32% | 1.97E-05 | HMPREF1322_RS06465 | - | hypothetical protein | Yes | HMPREF1322_RS06460,HMPREF1322_RS06465,<br>HMPREF1322_RS06470 |
| NZ_AJZS01000013.1 | 2448 | - | 11.90% | 17.41% | 5.51% | 2.20E-05 | HMPREF1322_RS00790 | *gyrA* | DNA gyrase subunit A | Yes | HMPREF1322_RS00785,HMPREF1322_RS00790 |
| NZ_AJZS01000050.1 | 1769 | - | 49.12% | 2.56% | 46.56% | 2.20E-05 | - | - | - | - | HMPREF1322_RS04325,HMPREF1322_RS04330 |
| NZ_AJZS01000066.1 | 22943 | + | 70.43% | 31.90% | 38.53% | 2.21E-05 | HMPREF1322_RS06180 | - | Ppx/GppA family<br>phosphatase | No | HMPREF1322_RS06175,HMPREF1322_RS06180,<br>HMPREF1322_RS06185 |
| NZ_AJZS01000050.1 | 797 | - | 38.96% | 13.79% | 25.16% | 2.31E-05 | HMPREF1322_RS04325 | - | DUF2436 domain-containing protein | Yes | HMPREF1322_RS04325 |
| NZ_AJZS01000011.1 | 18653 | + | 4.65% | 18.69% | 14.04% | 2.34E-05 | HMPREF1322_RS00745 | - | ABC transporter ATP-binding protein | No | HMPREF1322_RS00740,HMPREF1322_RS00745 |
| NZ_AJZS01000009.1 | 3668 | + | 12.68% | 21.61% | 8.93% | 2.40E-05 | HMPREF1322_RS00555 | - | DEAD/DEAH box helicase family protein | No | HMPREF1322_RS00555,HMPREF1322_RS00560 |
| NZ_AJZS01000011.1 | 13430 | + | 9.65% | 38.76% | 29.11% | 2.55E-05 | HMPREF1322_RS00725 | - | lactate utilization protein | Yes | HMPREF1322_RS00720,HMPREF1322_RS00725 |
| NZ_AJZS01000050.1 | 2103 | - | 61.60% | 6.56% | 55.04% | 2.66E-05 | HMPREF1322_RS04330 | - | nucleoside permease | Yes | HMPREF1322_RS04325,HMPREF1322_RS04330 |

LiH: growth in limited hemin
ExH: growth in excess hemin

We explored evidence for putative DNA sequence motifs at the identified all-context DMAs and DMCs. First, we observed that the DMAs identified in this all-context analysis (Table 3) did not overlap with our ‘GATC’-DMM results (S5 Table). We observed, however, that the 4-alpha-glucanotransferase HMPREF1322_RS03650 and the DUF3298 domain-containing protein HMPREF1322_RS07770 contained respectively two and one DMAs within 500 bp of a differentially methylated ‘GATC’ motif. Overall, DMAs and ‘GATC’-DMMs shared 13 genes annotated within 1 kb of their location (out of 75 and 85 genes respectively identified for DMAs and ‘GATC’-DMMs) (S5 Table). Second, we searched for over-representation of up to 15-nucleotide long sequences surrounding all DMAs and DMCs. Although the analyses identified putative sequences of length 10-11 nucleotides, the results did not reach statistical significance (MEME E-value > 0.05; S5 Fig).

### Genes with coordinated differential DNA methylation and expression changes

We assessed whether the observed DEGs also exhibited DNA methylation changes for 6mA and 5mC. We first compared DEGs to DMAs and DMCs each (Fig 7), and then explored further genomic regions that contained DEGs, DMAs, and DMCs (Fig 7).

**Fig 7.**
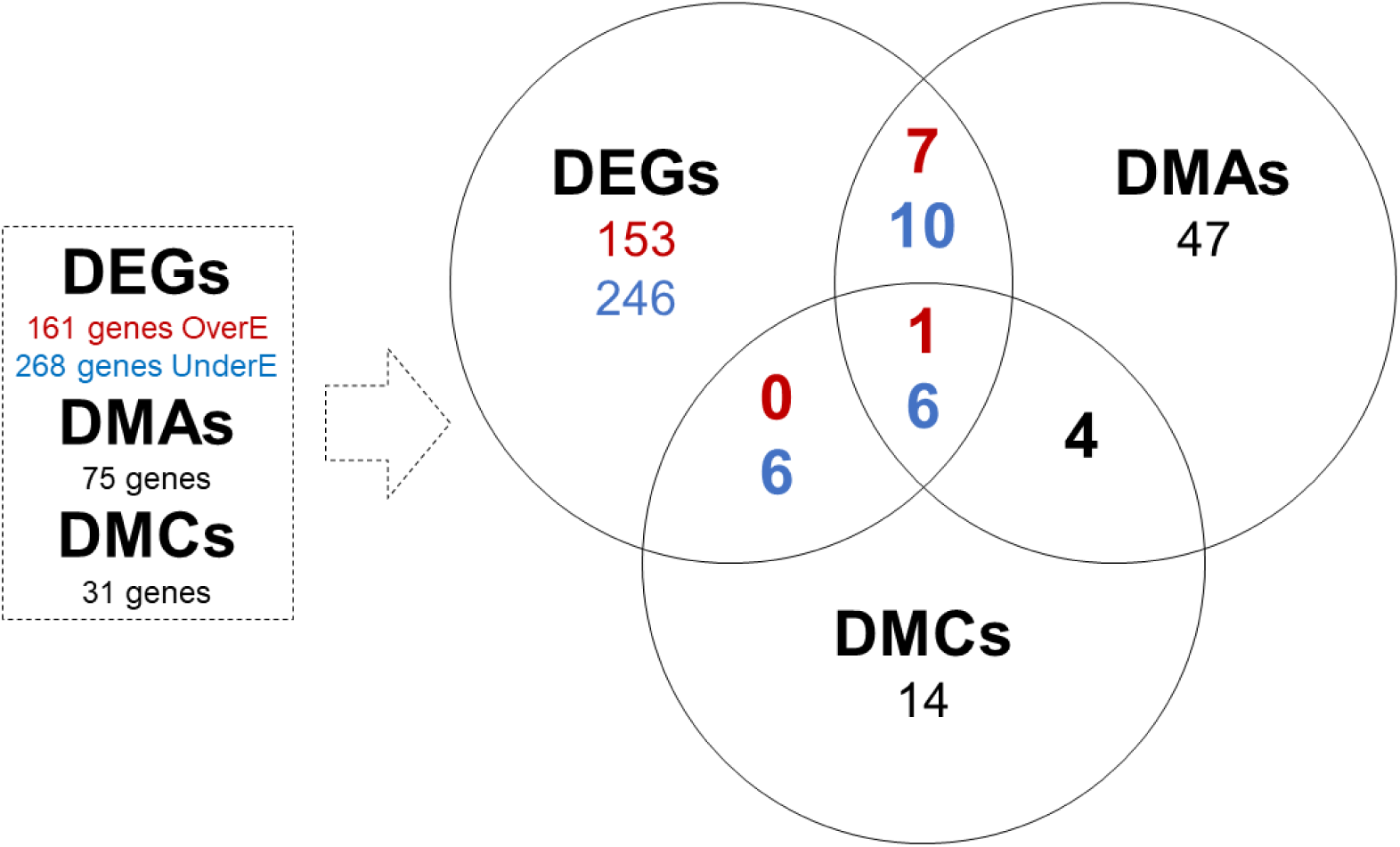
Overlap of genes with DEGs, DMAs and DMCs in excess hemin conditions.

Overall, 23 DEGs also harboured DMAs (Table 5; Table 6). Of these, 8 genes were over-expressed in excess hemin including the SDR family oxidoreductase HMPREF1322_RS07305 (*P. gingivalis* W83: PG2069) and the Ppx/GppA family phosphatase HMPREF1322_RS06180 (W83: PG1739) genes (Table 5). In all 8 cases the DMA was on the same strand as the gene, located either in or upstream of the gene (Table 5), with hypermethylation in excess hemin for most cases. The 15 DEGs with DMAs under-expressed in excess hemin included the hypothetical protein HMPREF1322_RS00730 (W83: PG1173) in the lactate utilization cluster, the lactate utilization protein HMPREF1322_RS00725 (W83: PG1172), the 4-alpha-glucanotransferase HMPREF1322_RS03650 (W83: PG0767) and ABC transporter protein gene cluster HMPREF1322_RS00745 (W83: PG1176) and HMPREF1322_RS00740 (W83: PG1175) (Table 6). Seven of the 15 genes had DMAs located upstream of the gene and six of these showed increased 6mA accompanied by decreased gene expression, indicating a potential regulatory effect of 6mA in dampening the expression of *P. gingivalis* genes according to hemin availability in the bacterium’s microenvironment. Most DMAs annotated to under-expressed genes had higher adenine methylation in excess hemin (84%) regardless of their location with respect to gene structure. In comparison to all-context DMAs, the ‘GATC’-DMMs were annotated to 14 DEGs (6 and 8 DEGs respectively over and under-expressed; S9 and S10 Tables). DEGs associated with ‘GATC’-DMMs included among others the 4-alpha-glucanotransferase HMPREF1322_RS03650 (W83: PG0767) mentioned immediately above. However, only four genes had ‘GATC’-DMMs upstream their start site, and another only two genes showed greater methylation associated with lower expression overall. This suggests very limited potential for a role of GATC methylation in the regulation of gene expression in *P. gingivalis* W50 in response to hemin availability.

**Table 5.** Genes over-expressed in excess hemin conditions with DMAs (> 1.5 LFC, > 5% methylation difference, FDR = 5%).

| Locus Tag | Gene Symbol | Gene Product | Contig | Gene start | Gene end | Strand Expr. | Methylated Position | Strand Meth. | Distance Meth. | DE analysis |  |  | DM analysis |  |  |  |
| --- | --- | --- | --- | --- | --- | --- | --- | --- | --- | --- | --- | --- | --- | --- | --- | --- |
|  |  |  |  |  |  |  |  |  |  | baseMean | LFC | P-value | mean % methylation: LiH | mean % methylation: ExH | % methylation difference | P-value |
| HMPREF1322_RS07305 | - | SDR family oxidoreductase | NZ_AJZS01000071.1 | 5169 | 5900 | - | 5863 | - | 694 | 39671.02 | 3.52 | 3.70E-20 | 62.88% | 80.42% | 17.54% | 6.37E-06 |
| HMPREF1322_RS07315 | *pdxA* | 4-hydroxythreonine-4-phosphate dehydrogenase PdxA | NZ_AJZS01000071.1 | 6386 | 7483 | - | 5863 | - | -523 | 10873.97 | 2.21 | 2.04E-19 | 62.88% | 80.42% | 17.54% | 6.37E-06 |
| HMPREF1322_RS06180 | - | Ppx/GppA family phosphatase | NZ_AJZS01000066.1 | 22201 | 23108 | - | 22242 | - | 41 | 914.83 | 1.59 | 2.19E-12 | 15.95% | 96.80% | 80.86% | 3.66E-07 |
| HMPREF1322_RS06180 | - | Ppx/GppA family phosphatase | NZ_AJZS01000066.1 | 22201 | 23108 | - | 22938 | - | 737 | 914.83 | 1.59 | 2.19E-12 | 85.99% | 37.41% | 48.59% | 4.56E-06 |
| HMPREF1322_RS01480 | - | tRNA-Thr | NZ_AJZS01000017.1 | 23424 | 23496 | - | 23369 | - | -55 | 362.80 | 2.29 | 2.35E-12 | 13.67% | 18.77% | 5.10% | 1.66E-05 |
| HMPREF1322_RS01485 | - | hypothetical protein | NZ_AJZS01000017.1 | 23557 | 24009 | - | 23369 | - | -188 | 2236.75 | 2.55 | 9.57E-12 | 13.67% | 18.77% | 5.10% | 1.66E-05 |
| HMPREF1322_RS04280 | *dnaE* | DNA polymerase III subunit alpha | NZ_AJZS01000048.1 | 883 | 4569 | - | 4837 | - | 3954 | 4976.97 | 1.52 | 1.59E-09 | 0.00% | 9.02% | 9.02% | 1.15E-05 |
| HMPREF1322_RS04100 | - | hypothetical protein | NZ_AJZS01000046.1 | 2632 | 4023 | - | 4018 | - | 1386 | 3442.27 | 1.59 | 9.47E-09 | 3.69% | 15.60% | 11.90% | 6.52E-06 |
| HMPREF1322_RS04100 | - | hypothetical protein | NZ_AJZS01000046.1 | 2632 | 4023 | - | 4605 | - | 1973 | 3442.27 | 1.59 | 9.47E-09 | 6.69% | 0.00% | 6.69% | 1.30E-05 |
| HMPREF1322_RS09370 | *rrf* | 5S ribosomal RNA | NZ_AJZS01000099.1 | 3390 | 3500 | - | 2869 | - | -521 | 5793.67 | 1.71 | 6.69E-04 | 55.68% | 47.10% | 8.58% | 1.41E-06 |

Strand Expr: Strand of the gene Strand Meth: Strand of the methylated position Distance Meth: Distance of the methylated position to the gene start site in base pairs, negative values indicate upstream location DE analysis: results from the differential expression analysis DM analysis: results from the differential methylation analysis LFC: log<sub>2</sub> fold change LiH: growth in limited hemin ExH: growth in excess hemin

**Table 6.**
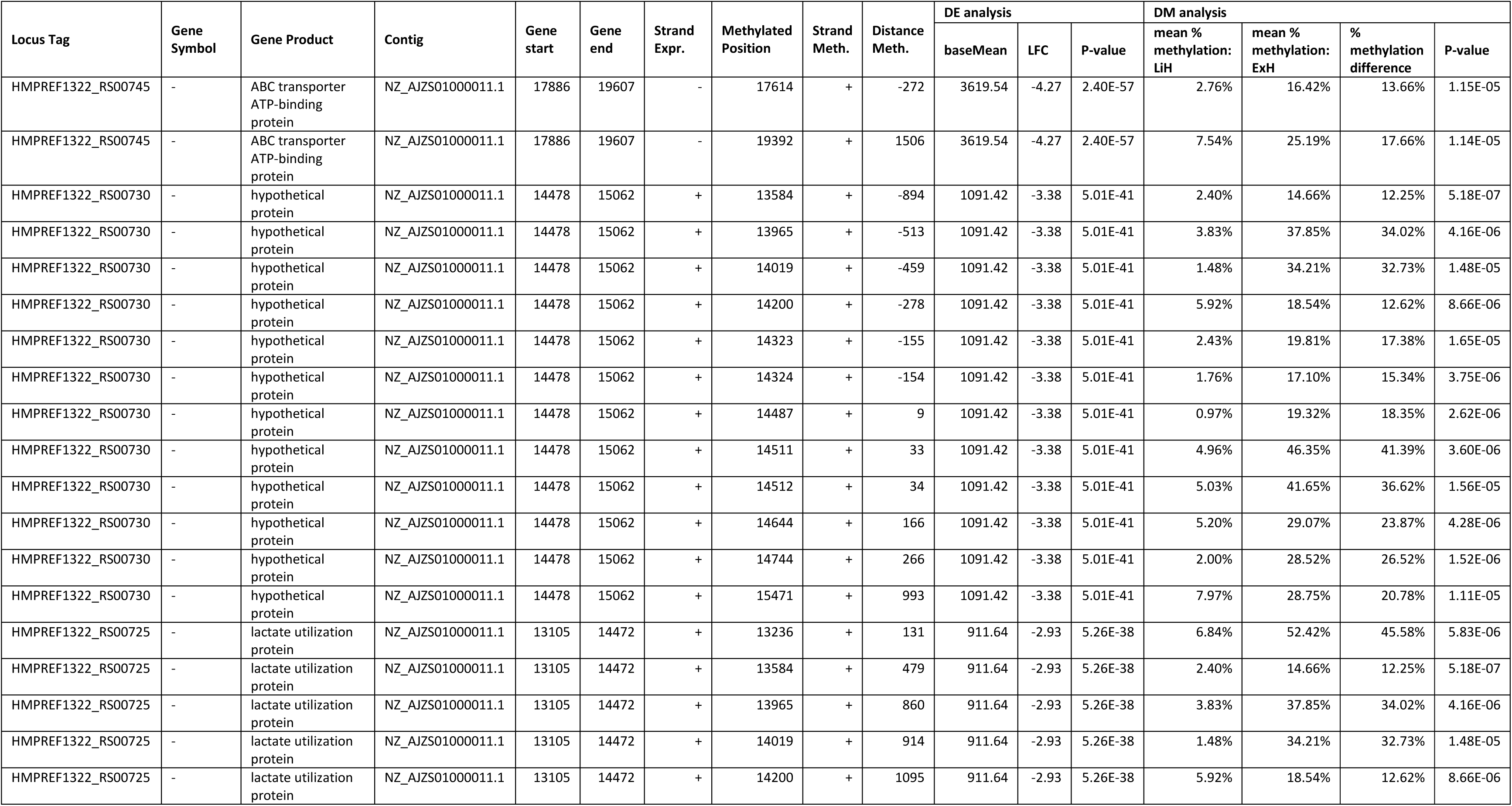
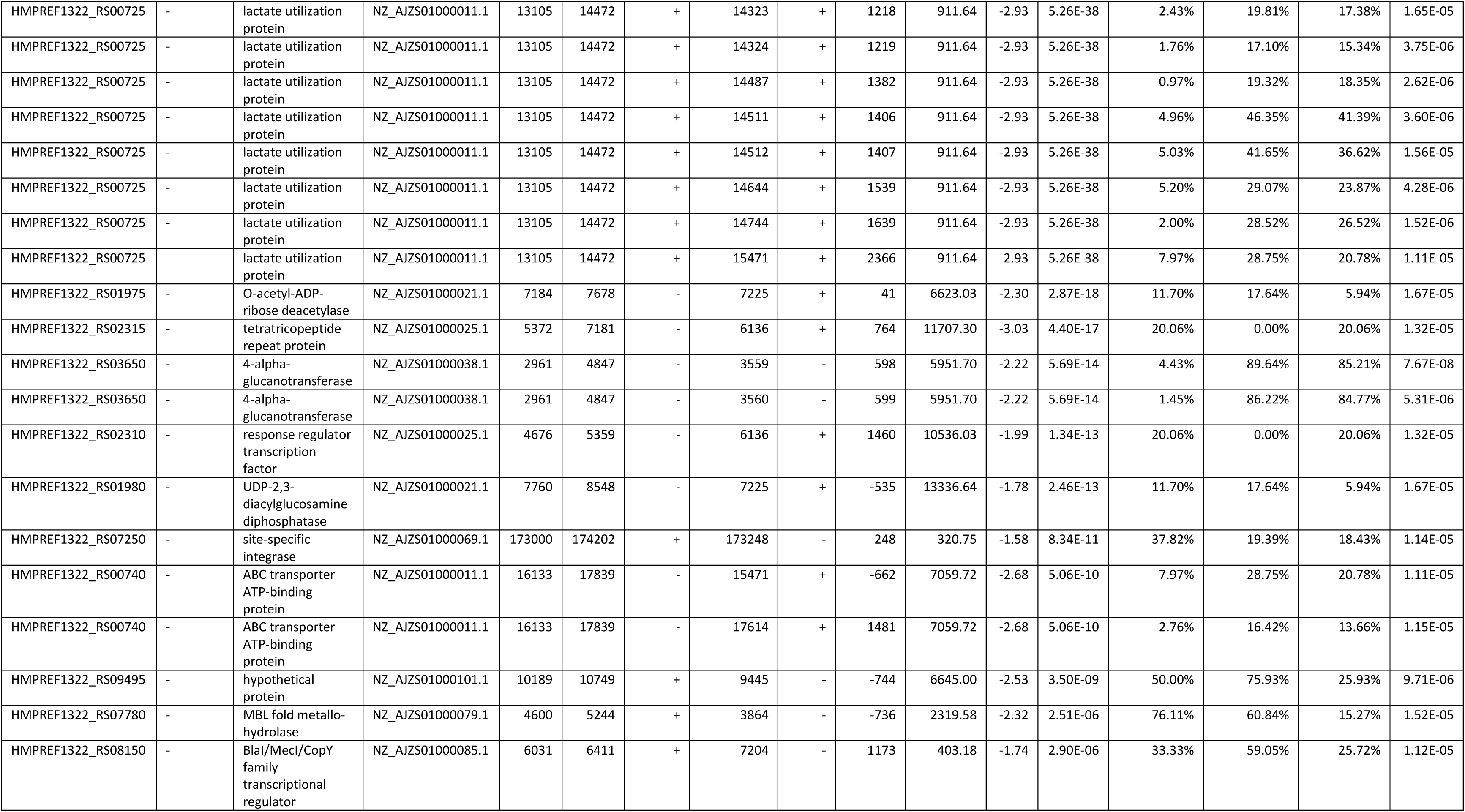
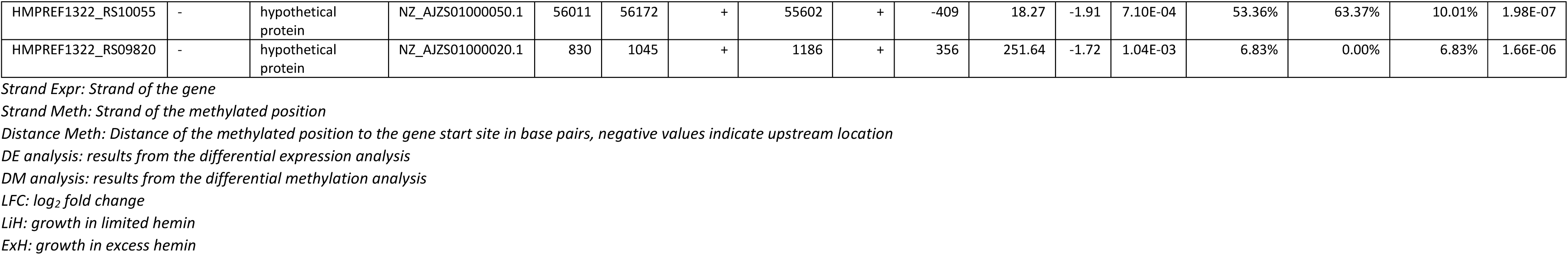
Genes under-expressed in excess hemin conditions with DMAs (< -1.5 LFC, > 5% methylation difference, FDR = 5%).

One over-expressed and 11 under-expressed genes also had DMCs within 1 kb of the gene (Tables 7 and 8). Two of the 11 under-expressed genes had DMCs upstream of the gene (Table 8) with increased methylation levels observed for all 11 genes. Moreover, 95% of DMCs annotated to under-expressed genes had increased cytosine methylation in excess hemin regardless of their location in relation to the gene. This is consistent with results from 6mA, although knowledge of gene regulation by 5mC methylation in bacteria remains limited.

**Table 7.** Genes over-expressed in excess hemin conditions with DMCs (> 1.5 LFC, > 5% methylation difference, FDR = 5%).

| Locus Tag | Gene Symbol | Gene Product | Contig | Gene start | Gene end | Strand Expr. | Methylated Position | Strand Meth. | Distance Meth. | DE analysis |  |  | DM analysis |  |  |  |
| --- | --- | --- | --- | --- | --- | --- | --- | --- | --- | --- | --- | --- | --- | --- | --- | --- |
|  |  |  |  |  |  |  |  |  |  | baseMean | LFC | P-value | mean % methylation: LiH | mean % methylation: ExH | % methylation difference | P-value |
| HMPREF1322_RS06180 | - | Ppx/GppA family phosphatase | NZ_AJZS01000066.1 | 22201 | 23108 | - | 22243 | - | 42 | 914.83 | 1.59 | 2.19E-12 | 10.21% | 64.13% | 53.91% | 6.11E-07 |
| HMPREF1322_RS06180 | - | Ppx/GppA family phosphatase | NZ_AJZS01000066.1 | 22201 | 23108 | - | 22943 | + | 742 | 914.83 | 1.59 | 2.19E-12 | 70.43% | 31.90% | 38.53% | 2.21E-05 |

Strand Expr: Strand of the gene Strand Meth: Strand of the methylated position Distance Meth: Distance of the methylated position to the gene start site in base pairs, negative values indicate upstream location DE analysis: results from the differential expression analysis DM analysis: results from the differential methylation analysis LFC: log<sub>2</sub> fold change LiH: growth in limited hemin ExH: growth in excess hemin

**Table 8.** Genes under-expressed in excess hemin conditions with DMCs (< -1.5 LFC, > 5% methylation difference, FDR = 5%).

| Locus Tag | Gene Symbol | Gene Product | Contig | Gene start | Gene end | Strand Expr. | Methylated Position | Strand Meth. | Distance Meth. | DE analysis |  |  | DM analysis |  |  |  |
| --- | --- | --- | --- | --- | --- | --- | --- | --- | --- | --- | --- | --- | --- | --- | --- | --- |
|  |  |  |  |  |  |  |  |  |  | baseMean | LFC | P-value | mean % methylation: LiH | mean % methylation: ExH | % methylation difference | P-value |
| HMPREF1322_RS00745 | - | ABC transporter ATP-binding protein | NZ_AJZS01000011.1 | 17886 | 19607 | - | 18653 | + | 767 | 3619.54 | -4.27 | 2.40E-57 | 4.65% | 18.69% | 14.04% | 2.34E-05 |
| HMPREF1322_RS00730 | - | hypothetical protein | NZ_AJZS01000011.1 | 14478 | 15062 | + | 13730 | + | -748 | 1091.42 | -3.38 | 5.01E-41 | 13.59% | 58.15% | 44.56% | 1.13E-05 |
| HMPREF1322_RS00730 | - | hypothetical protein | NZ_AJZS01000011.1 | 14478 | 15062 | + | 14106 | + | -372 | 1091.42 | -3.38 | 5.01E-41 | 16.77% | 47.36% | 30.59% | 7.45E-07 |
| HMPREF1322_RS00730 | - | hypothetical protein | NZ_AJZS01000011.1 | 14478 | 15062 | + | 14542 | + | 64 | 1091.42 | -3.38 | 5.01E-41 | 15.67% | 34.20% | 18.53% | 8.76E-06 |
| HMPREF1322_RS00730 | - | hypothetical protein | NZ_AJZS01000011.1 | 14478 | 15062 | + | 14669 | + | 191 | 1091.42 | -3.38 | 5.01E-41 | 7.23% | 19.84% | 12.61% | 1.28E-05 |
| HMPREF1322_RS00730 | - | hypothetical protein | NZ_AJZS01000011.1 | 14478 | 15062 | + | 14693 | + | 215 | 1091.42 | -3.38 | 5.01E-41 | 14.07% | 38.73% | 24.67% | 1.77E-05 |
| HMPREF1322_RS00730 | - | hypothetical protein | NZ_AJZS01000011.1 | 14478 | 15062 | + | 14698 | + | 220 | 1091.42 | -3.38 | 5.01E-41 | 26.82% | 44.78% | 17.96% | 7.90E-06 |
| HMPREF1322_RS00730 | - | hypothetical protein | NZ_AJZS01000011.1 | 14478 | 15062 | + | 14706 | + | 228 | 1091.42 | -3.38 | 5.01E-41 | 2.91% | 21.70% | 18.79% | 1.52E-05 |
| HMPREF1322_RS00725 | - | lactate utilization protein | NZ_AJZS01000011.1 | 13105 | 14472 | + | 12945 | + | -160 | 911.64 | -2.93 | 5.26E-38 | 5.18% | 61.79% | 56.61% | 2.71E-06 |
| HMPREF1322_RS00725 | - | lactate utilization protein | NZ_AJZS01000011.1 | 13105 | 14472 | + | 12946 | + | -159 | 911.64 | -2.93 | 5.26E-38 | 4.71% | 58.08% | 53.37% | 2.09E-06 |
| HMPREF1322_RS00725 | - | lactate utilization protein | NZ_AJZS01000011.1 | 13105 | 14472 | + | 13002 | + | -103 | 911.64 | -2.93 | 5.26E-38 | 3.76% | 55.38% | 51.62% | 3.63E-06 |
| HMPREF1322_RS00725 | - | lactate utilization protein | NZ_AJZS01000011.1 | 13105 | 14472 | + | 13071 | + | -34 | 911.64 | -2.93 | 5.26E-38 | 4.20% | 21.99% | 17.79% | 4.48E-06 |
| HMPREF1322_RS00725 | - | lactate utilization protein | NZ_AJZS01000011.1 | 13105 | 14472 | + | 13178 | + | 73 | 911.64 | -2.93 | 5.26E-38 | 7.08% | 60.16% | 53.08% | 2.24E-06 |
| HMPREF1322_RS00725 | - | lactate utilization protein | NZ_AJZS01000011.1 | 13105 | 14472 | + | 13240 | + | 135 | 911.64 | -2.93 | 5.26E-38 | 8.76% | 43.87% | 35.12% | 1.61E-06 |
| HMPREF1322_RS00725 | - | lactate utilization protein | NZ_AJZS01000011.1 | 13105 | 14472 | + | 13430 | + | 325 | 911.64 | -2.93 | 5.26E-38 | 9.65% | 38.76% | 29.11% | 2.55E-05 |
| HMPREF1322_RS00725 | - | lactate utilization protein | NZ_AJZS01000011.1 | 13105 | 14472 | + | 13730 | + | 625 | 911.64 | -2.93 | 5.26E-38 | 13.59% | 58.15% | 44.56% | 1.13E-05 |
| HMPREF1322_RS00725 | - | lactate utilization protein | NZ_AJZS01000011.1 | 13105 | 14472 | + | 14106 | + | 1001 | 911.64 | -2.93 | 5.26E-38 | 16.77% | 47.36% | 30.59% | 7.45E-07 |
| HMPREF1322_RS00725 | - | lactate utilization protein | NZ_AJZS01000011.1 | 13105 | 14472 | + | 14542 | + | 1437 | 911.64 | -2.93 | 5.26E-38 | 15.67% | 34.20% | 18.53% | 8.76E-06 |
| HMPREF1322_RS00725 | - | lactate utilization protein | NZ_AJZS01000011.1 | 13105 | 14472 | + | 14669 | + | 1564 | 911.64 | -2.93 | 5.26E-38 | 7.23% | 19.84% | 12.61% | 1.28E-05 |
| HMPREF1322_RS00725 | - | lactate utilization protein | NZ_AJZS01000011.1 | 13105 | 14472 | + | 14693 | + | 1588 | 911.64 | -2.93 | 5.26E-38 | 14.07% | 38.73% | 24.67% | 1.77E-05 |
| HMPREF1322_RS00725 | - | lactate utilization protein | NZ_AJZS01000011.1 | 13105 | 14472 | + | 14698 | + | 1593 | 911.64 | -2.93 | 5.26E-38 | 26.82% | 44.78% | 17.96% | 7.90E-06 |
| HMPREF1322_RS00725 | - | lactate utilization protein | NZ_AJZS01000011.1 | 13105 | 14472 | + | 14706 | + | 1601 | 911.64 | -2.93 | 5.26E-38 | 2.91% | 21.70% | 18.79% | 1.52E-05 |
| HMPREF1322_RS00885 | - | U32 family peptidase | NZ_AJZS01000015.1 | 486 | 2393 | - | 952 | + | 466 | 3114.91 | -3.09 | 3.14E-27 | 13.70% | 22.02% | 8.33% | 6.03E-06 |
| HMPREF1322_RS00860 | *htpG* | molecular chaperone HtpG | NZ_AJZS01000014.1 | 5016 | 7070 | - | 5938 | + | 922 | 15163.58 | -2.87 | 2.96E-23 | 34.76% | 52.63% | 17.87% | 2.45E-06 |
| HMPREF1322_RS06465 | - | hypothetical protein | NZ_AJZS01000069.1 | 2552 | 3754 | - | 3434 | - | 882 | 153316.28 | -2.35 | 1.18E-19 | 51.80% | 41.48% | 10.32% | 1.97E-05 |
| HMPREF1322_RS03650 | - | 4-alpha-glucanotransferase | NZ_AJZS01000038.1 | 2961 | 4847 | - | 3558 | - | 597 | 5951.70 | -2.22 | 5.69E-14 | 5.53% | 95.22% | 89.69% | 4.16E-07 |
| HMPREF1322_RS06460 | - | transporter | NZ_AJZS01000069.1 | 784 | 2448 | - | 3434 | - | 2650 | 37619.48 | -2.01 | 1.02E-13 | 51.80% | 41.48% | 10.32% | 1.97E-05 |
| HMPREF1322_RS00740 | - | ABC transporter ATP-binding protein | NZ_AJZS01000011.1 | 16133 | 17839 | - | 18653 | + | 2520 | 7059.72 | -2.68 | 5.06E-10 | 4.65% | 18.69% | 14.04% | 2.34E-05 |
| HMPREF1322_RS03860 | - | histidinol phosphate phosphatase | NZ_AJZS01000041.1 | 3824 | 4342 | - | 4471 | - | 647 | 15144.78 | -1.52 | 1.09E-09 | 17.52% | 22.93% | 5.41% | 9.62E-06 |
| HMPREF1322_RS00880 | - | DUF1661 domain-containing protein | NZ_AJZS01000015.1 | 1 | 203 | - | 952 | + | 951 | 129.52 | -1.93 | 8.19E-07 | 13.70% | 22.02% | 8.33% | 6.03E-06 |

*Strand Expr: Strand of the gene* *Strand Meth: Strand of the methylated position* *Distance Meth: Distance of the methylated position to the gene start site in base pairs, negative values indicate upstream location* *DE analysis: results from the differential expression analysis* *DM analysis: results from the differential methylation analysis* *LFC: log<sub>2</sub> fold change* *LiH: growth in limited hemin* *ExH: growth in excess hemin*

Six DEGs had both DMAs and DMCs in their gene body or within ±1kb of the gene (Fig 7), and a subset of these DEGs clustered in a genomic region that includes the lactate utilization gene cluster (Fig 6). The 6 DEGs include the Ppx/GppA family phosphatase HMPREF1322_RS06180 (W83: PG1739), over-expressed in excess hemin, and the hypothetical protein HMPREF1322_RS00730 (W83: PG1173), lactate utilization protein HMPREF1322_RS00725 (W83: PG1172), 4-alpha-glucanotransferase HMPREF1322_RS03650 (W83: PG0767), ABC transporter protein HMPREF1322_RS00745 (W83: PG1176), and HMPREF1322_RS00740 (W83: PG1175), which were under-expressed in excess hemin (Table 9). We assessed whether genome-wide DEGs were enriched for DMAs and DMCs, but did not observe a significant enrichment overall (OR = 0.77 (95% CI: 0.45, 1.31), P-value = 0.38 and OR = 1.13 (95% CI: 0.49, 2.47), P-value = 0.85, respectively). However, we observed strong evidence for overall differences in upstream or gene body DNA methylation levels at specific DEGs overlapping DMAs and DMCs (S6 and S7 Fig).

**Table 9.** Differentially expressed genes with DMAs and DMCs in excess hemin conditions (< -1.5 LFC, > 5% methylation difference, FDR = 5%).

| Locus Tag | Gene Symbol | Gene Product | Contig | Gene start | Gene end | Strand Expr. | Methylated Position | Strand Meth. | Distance Meth. | DE analysis |  |  |  | DM analysis |  |  |  |  |
| --- | --- | --- | --- | --- | --- | --- | --- | --- | --- | --- | --- | --- | --- | --- | --- | --- | --- | --- |
|  |  |  |  |  |  |  |  |  |  | Direction | baseMean | LFC | P-value | Modification | mean % methylation: LiH | mean % methylation: ExH | % methylation difference | P-value |
| HMPREF1322_RS00725 | - | lactate utilization protein | NZ_AJZS01000011.1 | 13105 | 14472 | + | 12945 | + | -160 | Under expr. | 911.64 | -2.93 | 5.26E-38 | DMC | 5.18% | 61.79% | 56.61% | 2.71E-06 |
| HMPREF1322_RS00725 | - | lactate utilization protein | NZ_AJZS01000011.1 | 13105 | 14472 | + | 12946 | + | -159 | Under expr. | 911.64 | -2.93 | 5.26E-38 | DMC | 4.71% | 58.08% | 53.37% | 2.09E-06 |
| HMPREF1322_RS00725 | - | lactate utilization protein | NZ_AJZS01000011.1 | 13105 | 14472 | + | 13002 | + | -103 | Under expr. | 911.64 | -2.93 | 5.26E-38 | DMC | 3.76% | 55.38% | 51.62% | 3.63E-06 |
| HMPREF1322_RS00725 | - | lactate utilization protein | NZ_AJZS01000011.1 | 13105 | 14472 | + | 13071 | + | -34 | Under expr. | 911.64 | -2.93 | 5.26E-38 | DMC | 4.20% | 21.99% | 17.79% | 4.48E-06 |
| HMPREF1322_RS00725 | - | lactate utilization protein | NZ_AJZS01000011.1 | 13105 | 14472 | + | 13178 | + | 73 | Under expr. | 911.64 | -2.93 | 5.26E-38 | DMC | 7.08% | 60.16% | 53.08% | 2.24E-06 |
| HMPREF1322_RS00725 | - | lactate utilization protein | NZ_AJZS01000011.1 | 13105 | 14472 | + | 13236 | + | 131 | Under expr. | 911.64 | -2.93 | 5.26E-38 | DMA | 6.84% | 52.42% | 45.58% | 5.83E-06 |
| HMPREF1322_RS00725 | - | lactate utilization protein | NZ_AJZS01000011.1 | 13105 | 14472 | + | 13240 | + | 135 | Under expr. | 911.64 | -2.93 | 5.26E-38 | DMC | 8.76% | 43.87% | 35.12% | 1.61E-06 |
| HMPREF1322_RS00725 | - | lactate utilization protein | NZ_AJZS01000011.1 | 13105 | 14472 | + | 13430 | + | 325 | Under expr. | 911.64 | -2.93 | 5.26E-38 | DMC | 9.65% | 38.76% | 29.11% | 2.55E-05 |
| HMPREF1322_RS00725 | - | lactate utilization protein | NZ_AJZS01000011.1 | 13105 | 14472 | + | 13584 | + | 479 | Under expr. | 911.64 | -2.93 | 5.26E-38 | DMA | 2.40% | 14.66% | 12.25% | 5.18E-07 |
| HMPREF1322_RS00725 | - | lactate utilization protein | NZ_AJZS01000011.1 | 13105 | 14472 | + | 13730 | + | 625 | Under expr. | 911.64 | -2.93 | 5.26E-38 | DMC | 13.59% | 58.15% | 44.56% | 1.13E-05 |
| HMPREF1322_RS00725 | - | lactate utilization protein | NZ_AJZS01000011.1 | 13105 | 14472 | + | 13965 | + | 860 | Under expr. | 911.64 | -2.93 | 5.26E-38 | DMA | 3.83% | 37.85% | 34.02% | 4.16E-06 |
| HMPREF1322_RS00725 | - | lactate utilization protein | NZ_AJZS01000011.1 | 13105 | 14472 | + | 14019 | + | 914 | Under expr. | 911.64 | -2.93 | 5.26E-38 | DMA | 1.48% | 34.21% | 32.73% | 1.48E-05 |
| HMPREF1322_RS00725 | - | lactate utilization protein | NZ_AJZS01000011.1 | 13105 | 14472 | + | 14106 | + | 1001 | Under expr. | 911.64 | -2.93 | 5.26E-38 | DMC | 16.77% | 47.36% | 30.59% | 7.45E-07 |
| HMPREF1322_RS00725 | - | lactate utilization protein | NZ_AJZS01000011.1 | 13105 | 14472 | + | 14200 | + | 1095 | Under expr. | 911.64 | -2.93 | 5.26E-38 | DMA | 5.92% | 18.54% | 12.62% | 8.66E-06 |
| HMPREF1322_RS00725 | - | lactate utilization protein | NZ_AJZS01000011.1 | 13105 | 14472 | + | 14323 | + | 1218 | Under expr. | 911.64 | -2.93 | 5.26E-38 | DMA | 2.43% | 19.81% | 17.38% | 1.65E-05 |
| HMPREF1322_RS00725 | - | lactate utilization protein | NZ_AJZS01000011.1 | 13105 | 14472 | + | 14324 | + | 1219 | Under expr. | 911.64 | -2.93 | 5.26E-38 | DMA | 1.76% | 17.10% | 15.34% | 3.75E-06 |
| HMPREF1322_RS00725 | - | lactate utilization protein | NZ_AJZS01000011.1 | 13105 | 14472 | + | 14487 | + | 1382 | Under expr. | 911.64 | -2.93 | 5.26E-38 | DMA | 0.97% | 19.32% | 18.35% | 2.62E-06 |
| HMPREF1322_RS00725 | - | lactate utilization protein | NZ_AJZS01000011.1 | 13105 | 14472 | + | 14511 | + | 1406 | Under expr. | 911.64 | -2.93 | 5.26E-38 | DMA | 4.96% | 46.35% | 41.39% | 3.60E-06 |
| HMPREF1322_RS00725 | - | lactate utilization protein | NZ_AJZS01000011.1 | 13105 | 14472 | + | 14512 | + | 1407 | Under expr. | 911.64 | -2.93 | 5.26E-38 | DMA | 5.03% | 41.65% | 36.62% | 1.56E-05 |
| HMPREF1322_RS00725 | - | lactate utilization protein | NZ_AJZS01000011.1 | 13105 | 14472 | + | 14542 | + | 1437 | Under expr. | 911.64 | -2.93 | 5.26E-38 | DMC | 15.67% | 34.20% | 18.53% | 8.76E-06 |
| HMPREF1322_RS00725 | - | lactate utilization protein | NZ_AJZS01000011.1 | 13105 | 14472 | + | 14644 | + | 1539 | Under expr. | 911.64 | -2.93 | 5.26E-38 | DMA | 5.20% | 29.07% | 23.87% | 4.28E-06 |
| HMPREF1322_RS00725 | - | lactate utilization protein | NZ_AJZS01000011.1 | 13105 | 14472 | + | 14669 | + | 1564 | Under expr. | 911.64 | -2.93 | 5.26E-38 | DMC | 7.23% | 19.84% | 12.61% | 1.28E-05 |
| HMPREF1322_RS00725 | - | lactate utilization protein | NZ_AJZS01000011.1 | 13105 | 14472 | + | 14693 | + | 1588 | Under expr. | 911.64 | -2.93 | 5.26E-38 | DMC | 14.07% | 38.73% | 24.67% | 1.77E-05 |
| HMPREF1322_RS00725 | - | lactate utilization protein | NZ_AJZS01000011.1 | 13105 | 14472 | + | 14698 | + | 1593 | Under expr. | 911.64 | -2.93 | 5.26E-38 | DMC | 26.82% | 44.78% | 17.96% | 7.90E-06 |
| HMPREF1322_RS00725 | - | lactate utilization protein | NZ_AJZS01000011.1 | 13105 | 14472 | + | 14706 | + | 1601 | Under expr. | 911.64 | -2.93 | 5.26E-38 | DMC | 2.91% | 21.70% | 18.79% | 1.52E-05 |
| HMPREF1322_RS00725 | - | lactate utilization protein | NZ_AJZS01000011.1 | 13105 | 14472 | + | 14744 | + | 1639 | Under expr. | 911.64 | -2.93 | 5.26E-38 | DMA | 2.00% | 28.52% | 26.52% | 1.52E-06 |
| HMPREF1322_RS00725 | - | lactate utilization protein | NZ_AJZS01000011.1 | 13105 | 14472 | + | 15471 | + | 2366 | Under expr. | 911.64 | -2.93 | 5.26E-38 | DMA | 7.97% | 28.75% | 20.78% | 1.11E-05 |
| HMPREF1322_RS00730 | - | hypothetical protein | NZ_AJZS01000011.1 | 14478 | 15062 | + | 13584 | + | -894 | Under expr. | 1091.42 | -3.38 | 5.01E-41 | DMA | 2.40% | 14.66% | 12.25% | 5.18E-07 |
| HMPREF1322_RS00730 | - | hypothetical protein | NZ_AJZS01000011.1 | 14478 | 15062 | + | 13730 | + | -748 | Under expr. | 1091.42 | -3.38 | 5.01E-41 | DMC | 13.59% | 58.15% | 44.56% | 1.13E-05 |
| HMPREF1322_RS00730 | - | hypothetical protein | NZ_AJZS01000011.1 | 14478 | 15062 | + | 13965 | + | -513 | Under expr. | 1091.42 | -3.38 | 5.01E-41 | DMA | 3.83% | 37.85% | 34.02% | 4.16E-06 |
| HMPREF1322_RS00730 | - | hypothetical protein | NZ_AJZS01000011.1 | 14478 | 15062 | + | 14019 | + | -459 | Under expr. | 1091.42 | -3.38 | 5.01E-41 | DMA | 1.48% | 34.21% | 32.73% | 1.48E-05 |
| HMPREF1322_RS00730 | - | hypothetical protein | NZ_AJZS01000011.1 | 14478 | 15062 | + | 14106 | + | -372 | Under expr. | 1091.42 | -3.38 | 5.01E-41 | DMC | 16.77% | 47.36% | 30.59% | 7.45E-07 |
| HMPREF1322_RS00730 | - | hypothetical protein | NZ_AJZS01000011.1 | 14478 | 15062 | + | 14200 | + | -278 | Under expr. | 1091.42 | -3.38 | 5.01E-41 | DMA | 5.92% | 18.54% | 12.62% | 8.66E-06 |
| HMPREF1322_RS00730 | - | hypothetical protein | NZ_AJZS01000011.1 | 14478 | 15062 | + | 14323 | + | -155 | Under expr. | 1091.42 | -3.38 | 5.01E-41 | DMA | 2.43% | 19.81% | 17.38% | 1.65E-05 |
| HMPREF1322_RS00730 | - | hypothetical protein | NZ_AJZS01000011.1 | 14478 | 15062 | + | 14324 | + | -154 | Under expr. | 1091.42 | -3.38 | 5.01E-41 | DMA | 1.76% | 17.10% | 15.34% | 3.75E-06 |
| HMPREF1322_RS00730 | - | hypothetical protein | NZ_AJZS01000011.1 | 14478 | 15062 | + | 14487 | + | 9 | Under expr. | 1091.42 | -3.38 | 5.01E-41 | DMA | 0.97% | 19.32% | 18.35% | 2.62E-06 |
| HMPREF1322_RS00730 | - | hypothetical protein | NZ_AJZS01000011.1 | 14478 | 15062 | + | 14511 | + | 33 | Under expr. | 1091.42 | -3.38 | 5.01E-41 | DMA | 4.96% | 46.35% | 41.39% | 3.60E-06 |
| HMPREF1322_RS00730 | - | hypothetical protein | NZ_AJZS01000011.1 | 14478 | 15062 | + | 14512 | + | 34 | Under expr. | 1091.42 | -3.38 | 5.01E-41 | DMA | 5.03% | 41.65% | 36.62% | 1.56E-05 |
| HMPREF1322_RS00730 | - | hypothetical protein | NZ_AJZS01000011.1 | 14478 | 15062 | + | 14542 | + | 64 | Under expr. | 1091.42 | -3.38 | 5.01E-41 | DMC | 15.67% | 34.20% | 18.53% | 8.76E-06 |
| HMPREF1322_RS00730 | - | hypothetical protein | NZ_AJZS01000011.1 | 14478 | 15062 | + | 14644 | + | 166 | Under expr. | 1091.42 | -3.38 | 5.01E-41 | DMA | 5.20% | 29.07% | 23.87% | 4.28E-06 |
| HMPREF1322_RS00730 | - | hypothetical protein | NZ_AJZS01000011.1 | 14478 | 15062 | + | 14669 | + | 191 | Under expr. | 1091.42 | -3.38 | 5.01E-41 | DMC | 7.23% | 19.84% | 12.61% | 1.28E-05 |
| HMPREF1322_RS00730 | - | hypothetical protein | NZ_AJZS01000011.1 | 14478 | 15062 | + | 14693 | + | 215 | Under expr. | 1091.42 | -3.38 | 5.01E-41 | DMC | 14.07% | 38.73% | 24.67% | 1.77E-05 |
| HMPREF1322_RS00730 | - | hypothetical protein | NZ_AJZS01000011.1 | 14478 | 15062 | + | 14698 | + | 220 | Under expr. | 1091.42 | -3.38 | 5.01E-41 | DMC | 26.82% | 44.78% | 17.96% | 7.90E-06 |
| HMPREF1322_RS00730 | - | hypothetical protein | NZ_AJZS01000011.1 | 14478 | 15062 | + | 14706 | + | 228 | Under expr. | 1091.42 | -3.38 | 5.01E-41 | DMC | 2.91% | 21.70% | 18.79% | 1.52E-05 |
| HMPREF1322_RS00730 | - | hypothetical protein | NZ_AJZS01000011.1 | 14478 | 15062 | + | 14744 | + | 266 | Under expr. | 1091.42 | -3.38 | 5.01E-41 | DMA | 2.00% | 28.52% | 26.52% | 1.52E-06 |
| HMPREF1322_RS00730 | - | hypothetical protein | NZ_AJZS01000011.1 | 14478 | 15062 | + | 15471 | + | 993 | Under expr. | 1091.42 | -3.38 | 5.01E-41 | DMA | 7.97% | 28.75% | 20.78% | 1.11E-05 |
| HMPREF1322_RS00740 | - | ABC transporter ATP-binding protein | NZ_AJZS01000011.1 | 16133 | 17839 | - | 15471 | + | -662 | Under expr. | 7059.72 | -2.68 | 5.06E-10 | DMA | 7.97% | 28.75% | 20.78% | 1.11E-05 |
| HMPREF1322_RS00740 | - | ABC transporter ATP-binding protein | NZ_AJZS01000011.1 | 16133 | 17839 | - | 17614 | + | 1481 | Under expr. | 7059.72 | -2.68 | 5.06E-10 | DMA | 2.76% | 16.42% | 13.66% | 1.15E-05 |
| HMPREF1322_RS00740 | - | ABC transporter ATP-binding protein | NZ_AJZS01000011.1 | 16133 | 17839 | - | 18653 | + | 2520 | Under expr. | 7059.72 | -2.68 | 5.06E-10 | DMC | 4.65% | 18.69% | 14.04% | 2.34E-05 |
| HMPREF1322_RS00745 | - | ABC transporter ATP-binding protein | NZ_AJZS01000011.1 | 17886 | 19607 | - | 17614 | + | -272 | Under expr. | 3619.54 | -4.27 | 2.40E-57 | DMA | 2.76% | 16.42% | 13.66% | 1.15E-05 |
| HMPREF1322_RS00745 | - | ABC transporter ATP-binding protein | NZ_AJZS01000011.1 | 17886 | 19607 | - | 18653 | + | 767 | Under expr. | 3619.54 | -4.27 | 2.40E-57 | DMC | 4.65% | 18.69% | 14.04% | 2.34E-05 |
| HMPREF1322_RS00745 | - | ABC transporter ATP-binding protein | NZ_AJZS01000011.1 | 17886 | 19607 | - | 19392 | + | 1506 | Under expr. | 3619.54 | -4.27 | 2.40E-57 | DMA | 7.54% | 25.19% | 17.66% | 1.14E-05 |
| HMPREF1322_RS03650 | - | 4-alpha-glucanotransferase | NZ_AJZS01000038.1 | 2961 | 4847 | - | 3558 | - | 597 | Under expr. | 5951.7 | -2.22 | 5.69E-14 | DMC | 5.53% | 95.22% | 89.69% | 4.16E-07 |
| HMPREF1322_RS03650 | - | 4-alpha-glucanotransferase | NZ_AJZS01000038.1 | 2961 | 4847 | - | 3559 | - | 598 | Under expr. | 5951.7 | -2.22 | 5.69E-14 | DMA | 4.43% | 89.64% | 85.21% | 7.67E-08 |
| HMPREF1322_RS03650 | - | 4-alpha-glucanotransferase | NZ_AJZS01000038.1 | 2961 | 4847 | - | 3560 | - | 599 | Under expr. | 5951.7 | -2.22 | 5.69E-14 | DMA | 1.45% | 86.22% | 84.77% | 5.31E-06 |
| HMPREF1322_RS06180 | - | Ppx/GppA family phosphatase | NZ_AJZS01000066.1 | 22201 | 23108 | - | 22242 | - | 41 | Over expr. | 914.83 | 1.59 | 2.19E-12 | DMA | 15.95% | 96.80% | 80.86% | 3.66E-07 |
| HMPREF1322_RS06180 | - | Ppx/GppA family phosphatase | NZ_AJZS01000066.1 | 22201 | 23108 | - | 22243 | - | 42 | Over expr. | 914.83 | 1.59 | 2.19E-12 | DMC | 10.21% | 64.13% | 53.91% | 6.11E-07 |
| HMPREF1322_RS06180 | - | Ppx/GppA family phosphatase | NZ_AJZS01000066.1 | 22201 | 23108 | - | 22938 | - | 737 | Over expr. | 914.83 | 1.59 | 2.19E-12 | DMA | 85.99% | 37.41% | 48.59% | 4.56E-06 |
| HMPREF1322_RS06180 | - | Ppx/GppA family phosphatase | NZ_AJZS01000066.1 | 22201 | 23108 | - | 22943 | + | 742 | Over expr. | 914.83 | 1.59 | 2.19E-12 | DMC | 70.43% | 31.90% | 38.53% | 2.21E-05 |

Strand Expr: Strand of the gene
Strand Meth: Strand of the methylated position
Distance Meth: Distance of the methylated position to the gene start site in base pairs, negative values indicate upstream location
DE analysis: results from the differential expression analysis
DM analysis: results from the differential methylation analysis
LFC: log<sub>2</sub> fold change
LiH: growth in limited hemin
ExH: growth in excess hemin

For all DEGs that also harboured DMAs or DMCs, we analysed the overall levels of adenine (S6 Fig) and cytosine (S7 Fig) methylation upstream (−100 bp), downstream (+100 bp) and in the gene body. Differential adenine and cytosine methylation were observed across all regions analysed for the lactate utilization protein HMPREF1322_RS00725 (W83: PG1172) (upstream; gene body; downstream; Table 9; Fig 6). Differential adenine and cytosine methylation were also observed in the gene body of the ABC transporters HMPREF1322_RS00740 (W83: PG1175) and HMPREF1322_RS00745 (W83: PG1176), and hypothetical protein HMPREF1322_RS00730 (W83: PG1173) (S6 and S7 Figs). These ABC transporter proteins are located directly downstream of a lactate utilization gene cluster, which includes the lactate utilization protein and hypothetical protein, possibly highlighting a key region of the *P. gingivalis* genome regulated by DNA methylation (Fig 6). Overall changes in adenine and/or cytosine methylation were also for example observed in the body of the 4−alpha−glucanotransferase HMPREF1322_RS03650 (W83: PG0767) gene, and upstream the ABC transporter HMPREF1322_RS00740 (W83: PG1175) (S6 and S7 Figs).

## Discussion

We identified a significant impact of hemin availability in the growth medium on the *P. gingivalis* epigenome and transcriptome. Growth of *P. gingivalis* is restricted in the healthy gingival crevice due to a tight restriction by the host on hemin availability. However, during increasing inflammation in periodontal disease, the availability of heme-containing molecules is substantially increased. Hemin-dependent differentially modified signatures were identified for gene expression, as well as for 6mA and 5mC methylation profiles. Comparison of the signals highlighted a cluster of coordinated changes in genes related to lactate utilisation and in ABC transporters. The findings identify putative bacterial DNA methylation signatures that may play a regulatory role in the expression of genes, in response to hemin availability.

Many of the genes that were over-expressed in excess hemin encoded proteins that contain iron as part of their tertiary structures or iron binding domains in the polyprotein. Furthermore, Lys-gingipain (Kgp; S1 Table) was also over-expressed, and it is involved in iron acquisition and is essential for black pigment (μ-oxo bishaem) formation from hemin and its absence attenuates virulence (15). Members of the ABC transport complex (Fig 6B and Table 6), forming a classical operon, may potentially represent a novel hemin or alternative solute acquisition system that may influence the cellular phenotype. The enhanced expression of thiamine phosphate synthase is not only limited to this gene (S1 Table) of the pathway; the contig also encodes ThiH, ThiG, ThiC, ThiS and a hypothetical protein, all arranged in an apparent operon. Thus, the biosynthesis of the energy co-factor thiamine, vitamin B1, may constitute an important aspect of *P. gingivalis* metabolism. Rubrerythin offers protection of the anaerobe, *P. gingivalis*, against the toxicity of molecular oxygen and hydrogen peroxide; an isogenic mutant is labile under these conditions (26). In addition, there appears to be a preference for succinate/fumarate and α-ketoglutarate utilisation (S1 Table).

Many genes under-expressed in excess hemin were involved in hemin binding or transport. Some genes also encoded for DNA transfer functions such as TraC, TraE, TraF and TraJ. Interestingly, the gene encoding the largest protein component of the type IX secretion system (T9SS), Sov (SprA), and also genes encoding OmpH, PorN (GldN) and PorX along with five cargo proteins, are under-expressed in excess hemin (S2 Table). The complex transports over 30 cargo proteins, including known virulence factors, with a C-terminal domain (CTD) signal motif, to the external environment (27). The gene encoding the long fimbriae surface appendage, FimA, together with its accessory proteins of the *fim* operon are under-expressed in excess hemin. The genes encoding regulatory proteins (*e.g.* AraC, TetR (AcrR), and LuxR) follow a similar expression pattern. Many transposases were also under-expressed in excess hemin, whereas no transposases were over-expressed in excess hemin, suggesting that the expression of these mobile elements is a characteristic of a stress environment where access to iron is limited.

We compared our results with two previous studies exploring expression (12) and proteomic (13) differences in *P. gingivalis* according to hemin availability in the media. We observed that both Anaya-Bergman *et al.* (2015) and Veith *et al.* (2018) previously reported changes in the expression of the phosphomethylpyrimidine synthase ThiC, involved in thiamine biosynthesis, under excess hemin conditions. Moreover, both studies also reported under-expressed levels of the genes coding peroxiredoxin and a phosphotransferase. Separately, Anaya-Bergman *et al.* (2015) shared three over-expressed genes with our study, *i.e.* the 2-iminoacetate synthase thiH, thiazole synthase thiG and a thiamine phosphate synthase (W83: PG2109/HMPREF1322_RS02110), and Veith *et al.* (2018) results matched three of the over-expressed genes identified by us, *i.e.* the rubrerythrin family protein HMPREF1322_RS04040 (W83: PG0195) and two hypothetical proteins (W83: PG0717/HMPREF1322_RS01045 and PG2105/HMPREF1322_RS02130). Veith *et. al* (2018) proteomic results also identified proteins for 11 additional under-expressed genes found in our study. These genes coded the cobaltochelatase subunit CobN (W83: PG1553/HMPREF1322_RS06780), the heme-binding protein HmuY(W83: PG1551/HMPREF1322_RS06790), a histidinol phosphate phosphatase (W83: PG0555/HMPREF1322_RS03860), a MMPL family transporter (W83: PG1180/HMPREF1322_RS06005), a peroxiredoxin (W83: PG0618/HMPREF1322_RS06360), a phosphoribosyltransferase (W83: PG1513/HMPREF1322_RS06960), a phosphotransferase (W83: PG0456/HMPREF1322_RS00135), a T9SS type A sorting domain-containing protein (W83: PG0495/HMPREF1322_RS03170), a TonB-dependent receptor (W83: PG1552/HMPREF1322_RS06785), a transporter protein (W83: PG1626/HMPREF1322_RS06460) and three hypothetical proteins (W83: PG0350/HMPREF1322_RS02675, W83: PG0613/HMPREF1322_RS06380, W83: PG1555/HMPREF1322_RS06770). The HMPREF1322_RS06770 hypothetical protein, the HMPREF1322_RS06360 peroxiredoxin and the HMPREF1322_RS06460 transporter respectively matched the PG1555 MotA/TolQ/ExbB proton channel family, PG0618 alkyl peroxide reductase AhpC and PG1626 FADL outer membrane receptor protein annotated in *P. gingivalis* W83 strain genome.

Epigenetic mechanisms, including the methylation of specific DNA sequences by DNA methyltransferases, allow unicellular organisms to respond rapidly to environmental stresses (28). Here we showed that both adenines and cytosines can be differentially methylated in *P. gingivalis* according to hemin availability in its microenvironment. Unlike in eukaryotes, in bacteria 5mC is not the dominant DNA modification and instead exists alongside 4mC and 6mA, the latter being the most prevalent modification in prokaryotes (29). Here we characterized both 6mA and 5mC and verified higher levels of adenine methylation (mean = 23% and median = 13% for 6mA genome-wide) in comparison to cytosine methylation (mean = 18% and median = 13% for 5mC genome-wide). However, lack of previous DNA methylation studies in *P. gingivalis* and a general lack of knowledge of the most common genetic motifs targeted by DNA methyltransferases (DNMTs) in this bacterium limited the analyses that we could undertake, and may impact some of the interpretation of data.

We analysed DNA methylation changes in Dam/Dcm motifs (mean = 8% and median = 13% for Dam; mean = 0.02% and median = 0% for Dcm genome-wide) to investigate possible Dam and Dcm-like DNA methylation in *P. gingivalis*. However, we observed only limited evidence for Dam-like ‘GATC’ methylation effects that overlapped differential gene expression. Alternatively, and because no standard model for methylation calling in *P. gingivalis* exists, we used Nanopore reads and all-context predictive models in Tombo to estimate the proportion of reads harbouring methylated bases in our study. All-context models in turn introduced a higher degree of uncertainty in methylation calling in our analysis. Additionally, we used proportion of methylated reads as an estimate of methylation levels in each sample of bacterial cells. In our data most positions at which DNA methylation was detected were not fully methylated or demethylated, pointing to presence of heterogeneity in the level of DNA methylation across different bacterial cells from the same culture sample (S8 and S9 Fig).

DNA methylation has historically been associated with bacterial DNA restriction-modification systems, which protect bacterial cells from foreign DNAs (28). More recently DNA methylation has also been shown to play important roles in other aspects of bacterial biology such as timing of DNA replication, timing of transposition and conjugal transfer of plasmids, DNA repair and cell partitioning, and regulation of gene expression (28). Less is known however about how widespread the role of DNA methylation is in virulence and pathogenesis. Prior evidence suggests that the activity of DNMTs such as Dam can for example affect virulence in a number of pathogens including *Escherichia coli, Salmonella* spp., *Yersinia pseudotuberculosis* and *Vibrio cholerae* (30).

Here we showed that the availability of hemin – a critical micronutrient that modulates the expression of virulence factors in *P. gingivalis* – is linked to changes in the DNA methylome of this bacterium. We also showed that methylation changes were coordinated to alterations in the expression of lactate utilization and ABC transporter genes, however their exact role in *P. gingivalis* remains unknown. Previous research found that lactate utilization (in *Haemophilus influenzae and Staphylococcus aureus*) and ABC transport of iron, manganese and/or zinc ions across the membrane (*e.g.,* in *Yersinia pestis, Salmonella enterica* and *Streptococcus* spp.) is linked to increased virulence (31–33). In our study, differential methylation in lactate utilization and ABC transporter genes co-occurred with their decreased expression in excess hemin conditions. We hypothesize that the decreased expression of these genes in excess hemin, and conversely their relatively increased expression in limited hemin, could be related to *P. gingivalis* scavenging of nutrients according to their availability in the growth environment.

In conclusion, using Nanopore sequencing we were able to characterise for the first time the epigenetic landscape of *P. gingivalis* W50, and to compare epigenetic variation to changes in gene expression. We showed that iron availability in the bacterium’s microenvironment can lead to discrete DNA methylation changes potentially impacting the expression of genes related to growth and virulence in humans. There is, however, a need to better characterise the diversity of motifs targeted by different DNMTs, including from *P. gingivalis.* A more comprehensive characterisation of DNMTs and their targeted motifs in the bacterial world will allow better future exploring of DNA methylation in *P. gingivalis* with Oxford Nanopore.

## Materials and methods

### Continuous culturing conditions

#### Strain Culture

*P. gingivalis* W50 was cultured on Columbia blood agar (CBA; Oxoid) supplemented with 5% oxalated horse blood (Oxoid) incubated anaerobically (5% CO_2_/5% H_2_/90% N_2_) at 37°C for 72 hours. A single colony was inoculated into 100 mL of pre-reduced Brain Heart Infusion (BHI; Oxoid) broth supplemented with 5 mg L^-1^ filter sterilised (0.22 µM) hemin and incubated until late log phase (48-72 hours).

#### Continuous Culture Chemostat Configuration

A one litre, sealed glass chemostat was autoclave sterilised with 300 mL BHI *in situ*. 100 mL of *P. gingivalis* culture was inoculated into the chemostat through a designated inoculation tube, using a peristaltic pump (Watson Marlow 101U). Sterile BHI (supplemented with either 0.2 or 5 mg L^-1^ hemin for limited and excess conditions, respectively) was continuously fed at a flow rate of 20 mL h^-1^ to achieve the working volume of 700 mL, before being increased to 70 mL h^-1^, in conjunction with harvest/waste pump activation at the equivalent rate. These conditions set a fixed dilution rate of 0.1 h^-1^ which corresponds to constant mean generation time of 6.9 h. Steady state conditions were achieved after approximately 10 mean generations. Environmental conditions were controlled/monitored using an Applikon ADI 1030 Bio-controller and BioXpert V2 software (Applikon Biotechnology, The Netherlands). The pH of the culture was maintained automatically at pH7±0.1 using 1 M NaOH and 0.5 M HCl fed through a peristaltic pump. Temperature was preserved at 37 ± 0.3°C with a silicon heat jacket. The chemostat was continuously sparged with filter-sterilised 5% CO_2_/95% N_2_ to maintain anaerobic conditions and stirred at 40 rpm (Motor Controller ADI 1012, Applikon Biotechnology) (S10 Fig).

#### Experimental Design

Two independent chemostat experiments were performed, one fed with 5 mg L^-1^ hemin (excess) and one fed with 0.2 mg L^-1^ hemin (limitation). Post-inoculation, the optical density (OD_600_) of the culture was monitored every 12 hours using a benchtop spectrophotometer (Jenway 6305). Steady state was defined as three consecutive OD_600_ readings within 10% of each other. Total viable counts and culture purity were determined daily, following serial dilutions in phosphate buffered saline, inoculation of CBA plates in triplicates and growth in a Whitley A45 anaerobic Workstation (Don Whitley Scientific, UK). Sampling for DNA and RNA analysis was performed every 48 hours during steady state (**Error! Reference source not found.** 1). This represented biological replicates separated by five full volume replenishments of the chemostat vessel.

#### Nucleic Acid Sampling

During nucleic acid sampling periods the harvest bottle was replaced with a sterile collection bottle (on ice) and approximately 20 mL was collected. For DNA sampling, 1 mL aliquots were centrifuged at 13,000 rpm for 10 minutes (4°C), prior to supernatant removal and storage at −80°C. For RNA samples, ten 1 mL samples were similarly centrifuged, the supernatant decanted, and the pellet reconstituted in 1mL of DNA/RNA Shield (Zymo Research, USA), prior to storage at −80°C.

#### DNA extraction

One mL cultures were centrifuged at 17,000×g, at 4°C for 10 minutes. The pellets were used to extract high molecular weight DNA using Gentra Puregene yeast/bacteria DNA isolation kit (Qiagen), following the manufacturer’s protocol. DNA was finally resuspended in 200 µl EB buffer (Qiagen) and stored at 4°C before use.

#### RNA extraction

Cells were pelleted as above and resuspended/lysed in 350 µl of DNA/RNA Shield (Cambridge Biosciences) at room temperature. The solutions were used directly in RNA isolation with Zymo Quick-RNA™ miniprep kit (Cambridge Biosciences) in accordance with the manufacture’s protocol. This incorporated the initial chromosomal DNA removal, and on-column DNAase I digestion before the final wash and elution into 50 µl nuclease-free water. RNA was stored at −70°C.

#### Illumina RNA sequencing

RNA samples were first subjected to rRNA depletion using the NEBNext® rRNA Depletion Kit (Bacteria). The samples were then used for RNA-seq library preparation using the NEBNext Ultra II Directional RNA Library Prep Kit for Illumina. Both kits were used according to the manufacturer’s instructions. Sequencing was performed by Novagene, UK on an Illumina Novaseq6000 instrument.

#### Oxford Nanopore DNA sequencing

MinIon library preparation and DNA sequencing was performed at the University of Nottingham on MinIon flow cell (Oxford Nanopore). Whole-genome sequencing libraries were prepared from six bacterial genomic DNA samples. DNA samples were quantified using the Qubit 4 Fluorometer (Thermo Fisher Scientific) and the Qubit dsDNA BR Assay Kit (Thermo Fisher Scientific; Q32853) and the Agilent 4200 TapeStation and the Agilent Genomic DNA ScreenTape Assay (Agilent; 5067-5365 and 5067-5366) were used to assess the molecular weight of the DNA. Sequencing libraries were prepared using the Native barcoding genomic DNA protocol (Oxford Nanopore Technologies; Version: NBE_9065_v109_revS_14Aug2019). This protocol uses the Ligation Sequencing Kit (Oxford Nanopore Technologies; SQK-LSK109) and the Native Barcoding Expansion 1-12 kit (Oxford Nanopore Technologies; EXP-NBD104) to barcode the unsheared DNA fragments within each sample. Barcoded samples were then pooled in equimolar amounts for the final sequencing adapter ligation. All purification steps were performed using AMPure XP beads (Beckman Coulter; A63882). The final barcoded library pool was loaded onto a MinION flow cell (Oxford Nanopore Technologies; FLO-MIN106 R9.4.1) and run on the GridION X5 Mk1.

### Bioinformatic analyses

#### RNA sequencing data processing

Sequencing data was processed using the ProkSeq pipeline v2.8 (22) Briefly, FastQC (34)and AfterQC (35, 36) were implemented to check the quality of data, trim adaptor sequences and filter out low quality reads using standard parameters. This resulted in a total of ≈25.3–36.0 M PE reads (2 × 76 bp) being kept for downstream processing. Subsequently, bowtie2 (35) was used to align reads against the 104 contigs of the *P. gingivalis* W50 genome assembly deposited in the NCBI RefSeq (accession no. GCF_000271945.1). RefSeq gene annotations were extracted from *P. gingivalis*’ GTF file and the number of total reads per gene were calculated using featureCounts (37). Counts per gene were estimated for 1992 genes. RSeQC (38) was used to study sample coverage uniformity along the body of genes.

### DNA methylation calling

#### Motif-dependent DNA methylation calling

Modified basecalling was first performed with Guppy v3.2.10+aabd4ec (Oxford Nanopore) using the high accuracy methylation aware configuration available for Dam and Dcm motifs from *Escherichia coli* (‘dna_r9.4.1_450bps_modbases_dam-dcm-cpg_hac.cfg’). Total Nanopore read data included ≈240–380 K reads (88 bp – 143 kb, N50 = 27 kb) per sample. Methylated and canon bases were then aggregated against the *P. gingivalis* W50 reference genome using Medaka v0.11.5 (https://github.com/nanoporetech/medaka). Base counts were normalized for the total number of reads in each sample × 10,000 reads in similar fashion to the RPG10K method (39, 40). The proportion of methylated bases to the total number of bases (methylated + canon bases) was estimated at each genomic motif and motifs were filtered for minimum 10× coverage across samples. A total of 12,066 ‘GATC’ (Dam), 419 ‘CCAGG’ (Dcm) and 400 ‘CCTGG’ (Dcm) motifs were kept for downstream analyses.

#### Motif-independent DNA methylation calling

Nanopore signals were originally stored in multi-read fast5 files and the *ont_fast5_api* ‘multi_to_single_fast5’ (https://github.com/nanoporetech/ont_fast5_api) was used to generate single fast5 files before detection of non-standard nucleotides with Tombo v1.5.1 (41). First, electric current signal level data was re-squiggled using the same *P. gingivalis* W50 reference genome used to pre-process the RNA-Seq and methylated motif data. Secondly, all-context motif-independent models were used in Tombo to detect the alternative bases 6mA and 5mC from the signal data. Alternative base calling in each sample was summarized per genomic position with coverage and dampened fractions estimated at each adenine and cytosine site of both positive and negative DNA strands. Coverage was stored in the bedGraph format and proportion of methylation in the WIG format. Data files were imported into R using the package *rtracklayer* (42) and combined into a single data frame using GRanges, IRanges and *findOverlaps* from the IRanges package (43).

The proportion of methylation – dampened fraction – was estimated by adjusting the proportion of methylated reads according to coverage at low coverage sites in order to include them in downstream analyses (41). Despite this, only adenines and cytosines with at least 10× coverage across samples were included in our analysis here as well. Overall, 1,150,659 adenines and 1,079,809 cytosines were kept for downstream analyses, with 350–525× mean base coverage per sample.

### Statistical analyses

#### Differential gene expression analysis

DESeq2 (23) was used to normalize the read count data and perform differential gene expression analysis. Growth in limited hemin was the ‘control’ and growth in excess hemin was the ‘treatment’. All three samples from each condition were used and global sample differences were observed using the principal component analysis method. Both DESeq2 and edgeR (44) were implemented from ProkSeq’s downstream analysis. Both methods allow for inter-sample comparison in differential gene expression analysis. Here we used edgeR results as sensitivity for DESeq2 results since edgeR normalizes expression data differently to DESeq2 (DESeq2 uses the median of ratios method and edgeR uses the trimmed mean of M values to normalize read count data). The false discovery rate method (FDR) was used to identify differentially expressed genes and genes with FDR < 5% and a log_2_ fold change (LFC) > 1.5 or < −1.5 were respectively considered over and under-expressed in excess hemin. Differentially expressed genes were subsequently manually annotated to the *P. gingivalis* W83 genome (GenBank accession no. AE015924) (25) in order to compare results with previous research. Synteny analysis at specific loci was performed using MAUVE (45).

#### Gene ontology enrichment analysis

Gene ontology (GO) enrichment analysis in over and under-expressed gene sets was performed using ShinyGO (46). Because the genome of *P. gingivalis* W50 was not available from ShinyGO’s species database, only well-characterized genes that matched gene symbols in *P. gingivalis* W83 reference strain genome were used in this analysis. A more relaxed FDR threshold of 10% was applied to assess enrichment in this case and GO molecular functions were reported.

#### Differential DNA methylation analysis

Mean and median DNA methylation across samples were calculated using all motifs, adenines and cytosines kept for downstream analysis. Global DNA methylation differences between the samples were observed using principal component analysis for Dam/Dcm and all-context 6mA and 5mC modifications, selecting for minimum 10× and 100× coverage per position.

Two-sided Student’s t-tests were applied at each motif, adenine and cytosine site after filtering for positions where at least a 5% mean methylation difference was observed between the experimental conditions (79 ‘GATC’ motifs (Dam), 139,794 all-context adenines and 87,398 all-context cytosines). A sensitivity analysis was performed for minimum 100× coverage (66 ‘GATC’ motifs (Dam), 85,090 all-context adenines and 46,930 all-context cytosines) to ascertain whether the significance of results was impacted by depth of sequencing. Results were considered statistically significant at an FDR threshold of 5% to match gene expression analysis.

The differentially methylated motifs and all-context positions identified were annotated to nearby genes considering the location and gene strand. First, ‘GATC’-DMMs, DMAs and DMCs were annotated to the gene they were located if present in the gene body, and secondly, they were annotated to genes found in the same contig and within ±1kb of their location. Genes located in the opposite strand of the methylation signal were considered in both cases.

In comparing the overlap between DEGs, DMA, DMCs and ‘GATC’-DMMs we considered all DMAs, DMCs and ‘GATC’-DMMs that were in or near the gene (±1 kb away, both strands). DMA and DMC data were further used to test the wider DNA methylation effects on the expression of nearby genes. A contingency table containing all 1192 genes was constructed and Fischer’s exact tests were employed to test for the enrichment of genes with concurrent gene expression and all-context DNA methylation changes. The enrichment of differential DNA methylation directly upstream, downstream and in the body of genes was also tested. Here, Wilcoxon signed-rank tests were used to test for generalized shifts in DNA methylation signals between the experimental conditions 100 bp upstream, 100 bp downstream and within the body of genes previously annotated to DMAs and/or DMCs.

#### Motif enrichment analysis

The MEME (Multiple EM for Motif Elicitation) online tool (47) was used to detect enriched motifs containing the DMAs and DMCs identified with concurrent gene expression changes. For this, 15 base pair long DNA sequences surrounding the DMAs and DMCs identified were extracted. Subsequently, motifs with minimum 4 bp and maximum 15 bp were searched using MEME’s Classic objective function and the ZOOPS distribution model. Both the given strand and the reverse complement strand were considered, and the top 10 motifs found were reported. Motifs were considered significant if their reported E-value threshold was ≤ 0.05.

## Acknowledgments

We are grateful to the Medical Research Council of United Kingdom for the grant: MR/P012175/2. We thank the teams at the genome sequencing centres for assistance, and Mr Darioush Yarand and the Rosalind KCL high performance Linux cluster system administration team for help resolving file space and software installation queries.

## Author contributions

MAC and VR conceptualized the study. MAC, VR, DAD and JTB provided resources. MAC, VR, and JTB supervised the study. RC and JA-O implemented the methodology, JA-O performed experiments, and RC carried out formal analysis and software implementation. JJV, FR-A, SJ, DAD, PDM, and VR contributed to methodology, experiments, analysis, and results interpretation. RC, JA-O, MAC, and JTB drafted the manuscript, and all authors reviewed and edited the manuscript.

## Supporting information

**S1 Fig.** Overlap between DeSeq2 and edgeR results for differentially expressed *P. gingivalis* genes cultured in excess hemin conditions.

**S2 Fig.** DESeq2-normalized counts PCA for *P. gingivalis* growth in limited (red) and excess (blue) hemin conditions.

**S3 Fig.** All-context DNA methylation PCA for *P. gingivalis* growth in limited (LiH) and excess (ExH) hemin conditions, selecting for minimum 10× (A) and 100× (B) coverage.

**S4 Fig.** Dam/Dcm DNA methylation PCA for *P. gingivalis* growth in limited (LiH) and excess (ExH) hemin conditions, selecting for minimum 10× (A) and 100× (B) coverage.

**S5 Fig.** MEME motif analysis results for the 49 and 47 DMAs and DMCs identified in the main analysis.15-nucleotide long sequences surrounding DMAs and DMCs were extracted, and motifs were analysed considering minimum and maximum motif widths of 4 and 15 nucleotides, respectively. A maximum number of 10 motifs were searched. No motifs reached significance (E-value < 0.05).

**S6 Fig.** Adenine methylation patterns upstream (−100 bp), downstream (+100 bp) and in the gene body of differentially expressed genes annotated to DMAs. Differential methylation significance was tested using Wilcoxon signed-rank tests between limited (red) and excess (blue) hemin conditions.

**S7 Fig.** Cytosine methylation patterns upstream (−100 bp), downstream (+100 bp) and in the gene body of differentially expressed genes annotated to DMAs. Differential methylation significance was tested using Wilcoxon signed-rank tests between limited (red) and excess (blue) hemin conditions.

**S8 Fig.** Distribution of all-context DNA methylation for *P. gingivalis* growth in limited (LiH) and excess (ExH) hemin conditions, selecting for 10× (A) and 100× (B) coverage.

**S9 Fig.** Distribution of Dam/Dcm DNA methylation for *P. gingivalis* growth in limited (LiH) and excess (ExH) hemin conditions, selecting for 10× (A) and 100× (B) coverage.

**S10 Fig.** Schematic representation of chemostat set up in class II fume cabinet.

**S1 Table.** Genes over-expressed in excess hemin conditions (>1.5 LFC, FDR = 5%).

**S2 Table.** Genes under-expressed in excess hemin conditions (<−1.5 LFC, FDR = 5%).

**S3 Table.** GO molecular functions over-expressed in excess hemin conditions (based on 40 genes, FDR = 10%).

**S4 Table.** GO molecular functions under-expressed in excess hemin conditions (based on 8 genes, FDR = 10%).

**S5 Table.** ’GATC’-DMMs identified in *P. gingivalis* W50, selecting for minimum 10× coverage and mean 5% methylation difference between experimental conditions (FDR = 5%).

**S6 Table.** ’GATC’-DMMs identified in *P. gingivalis* W50, selecting for minimum 100× coverage and mean 5% methylation difference between experimental conditions (FDR = 5%).

**S7 Table.** DMAs (6mA) identified in *P. gingivalis* W50, selecting for minimum 100× coverage and mean 5% methylation difference between experimental conditions (FDR = 5%).

**S8 Table.** DMCs (5mC) identified in *P. gingivalis* W50, selecting for minimum 100× coverage and mean 5% methylation difference between experimental conditions (FDR = 5%).

**S9 Table.** Genes over-expressed in excess hemin conditions with ’GATC’-DMMs (> 1.5 LFC, > 5% methylation difference, FDR = 5%).

**S10 Table.** Genes under-expressed in excess hemin conditions with ’GATC’-DMMs (> 1.5 LFC, > 5% methylation difference, FDR = 5%).

## Notes

### Competing Interest Statement

The authors have declared no competing interest.

