## Supplemental Figures for "Hemin availability induces coordinated DNA methylation and gene expression changes in *Porphyromonas gingivalis*"

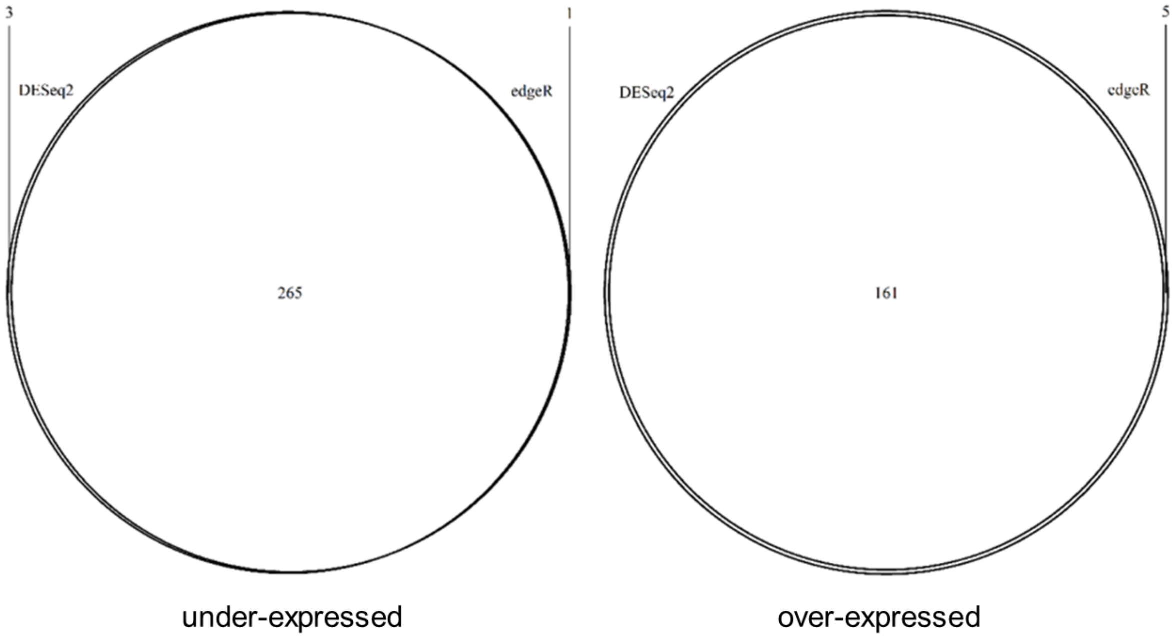


S1 Fig. Overlap between DESeq2 and edgeR results for differentially expressed *P. gingivalis* genes cultured in excess hemin conditions.

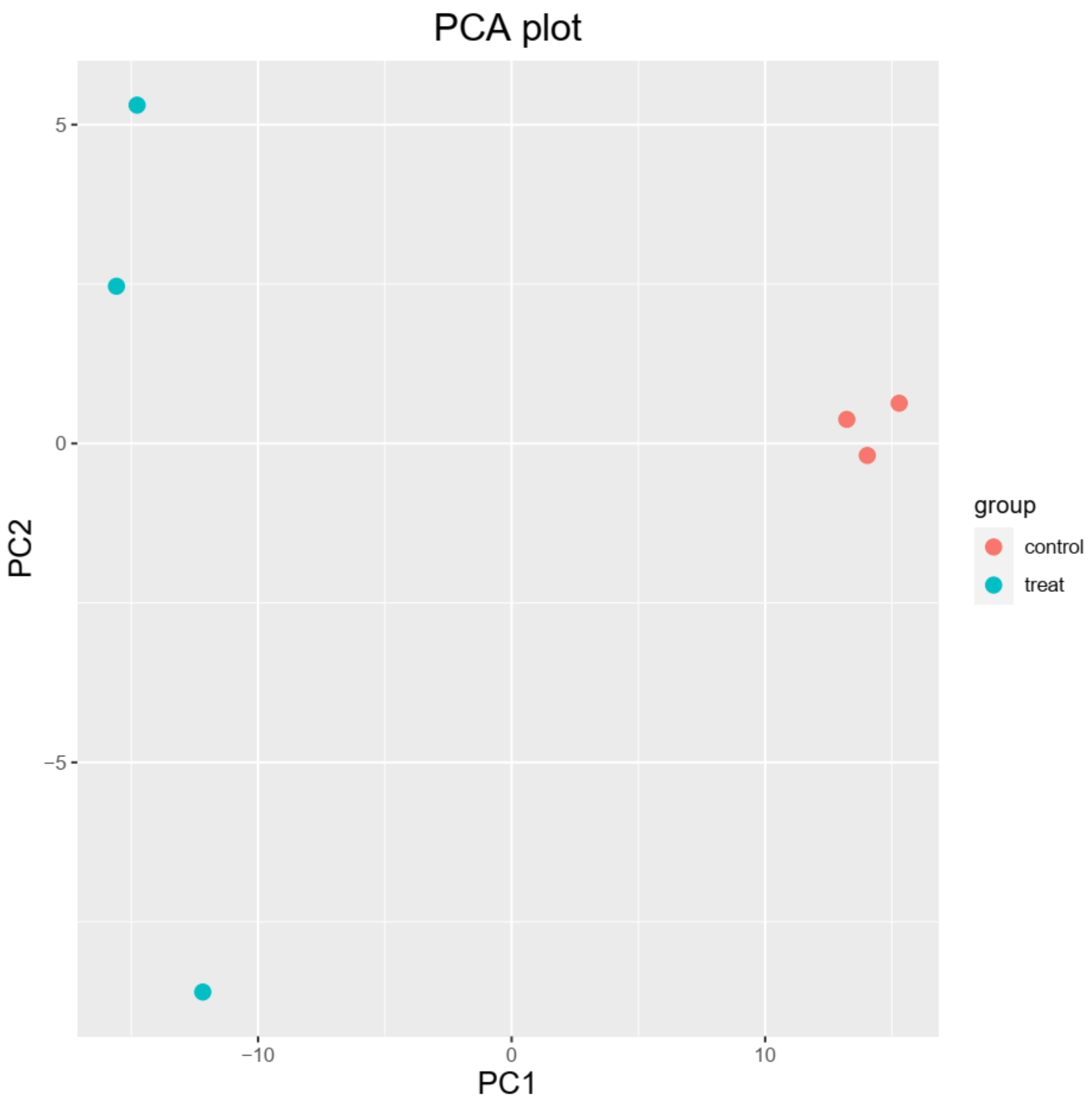


S2 Fig. DESeq2-normalized counts PCA for *P. gingivalis* growth in limited (red) and excess (blue) hemin conditions.

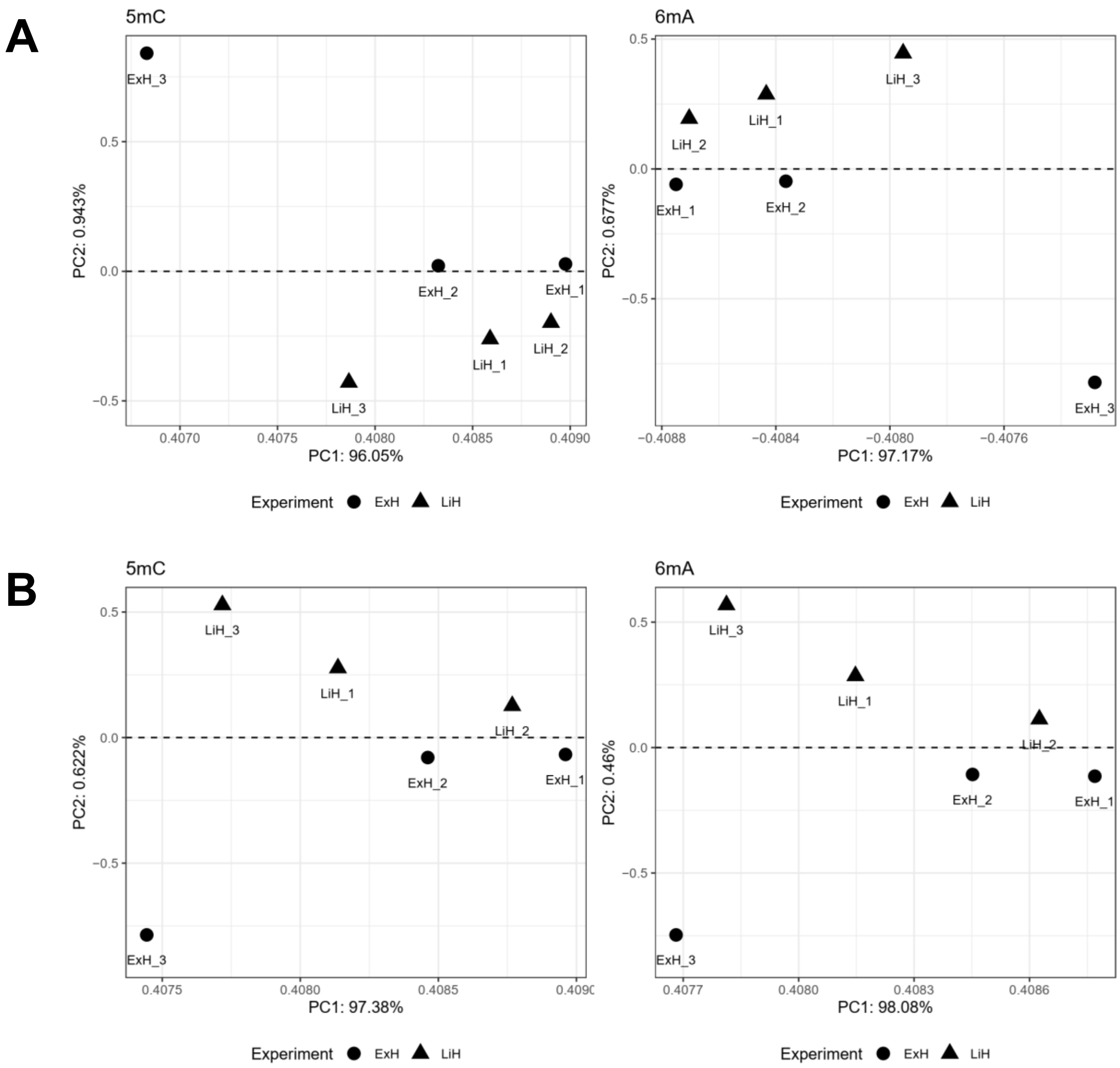


S3 Fig. All-context DNA methylation PCA for *P. gingivalis* growth in limited (LiH) and excess (ExH) hemin conditions, selecting for minimum 10× (A) and 100× (B) coverage.

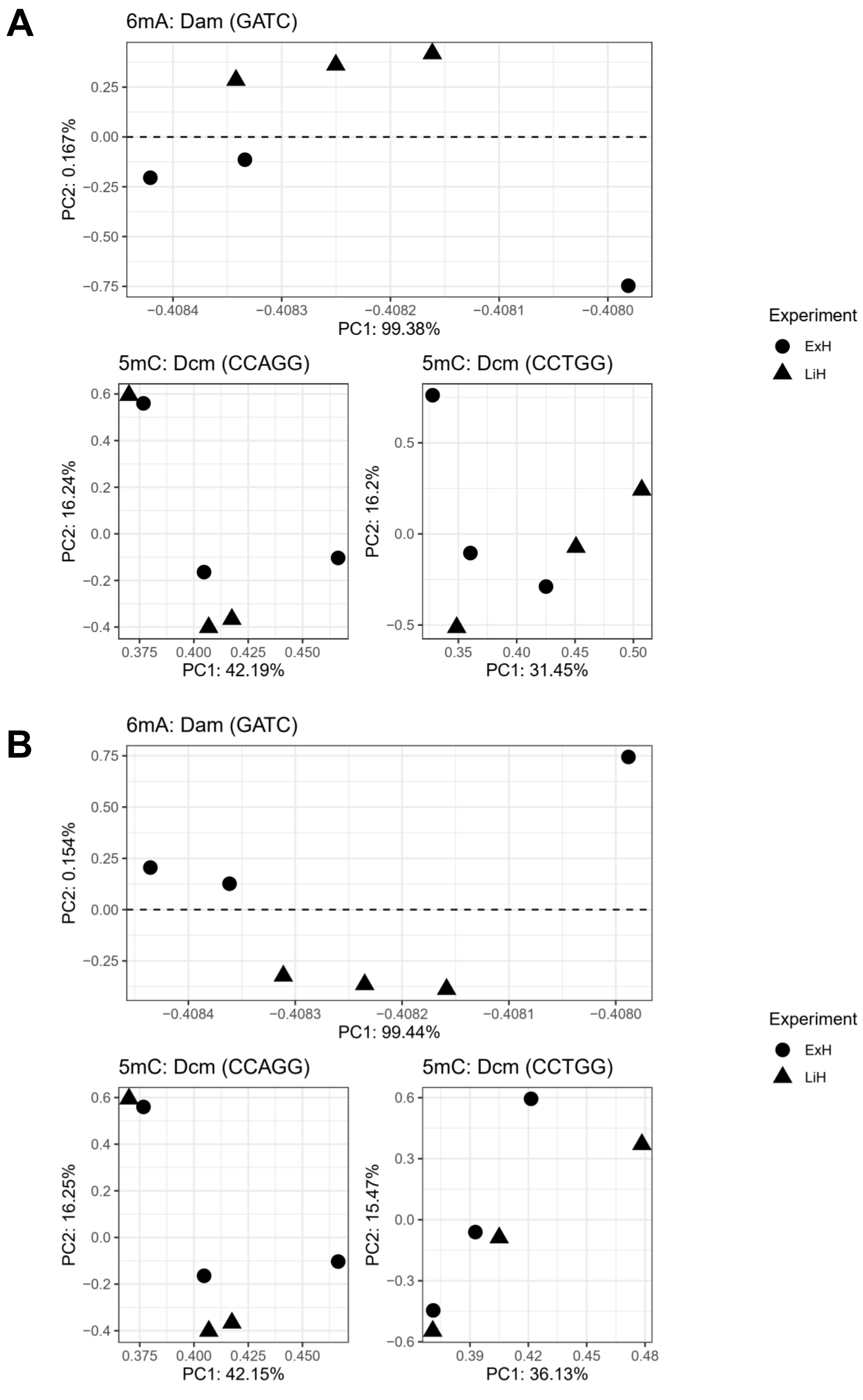


S4 Fig. Dam/Dcm DNA methylation PCA for *P. gingivalis* growth in limited (LiH) and excess (ExH) hemin conditions, selecting for minimum 10× (A) and 100× (B) coverage.

### DMAs

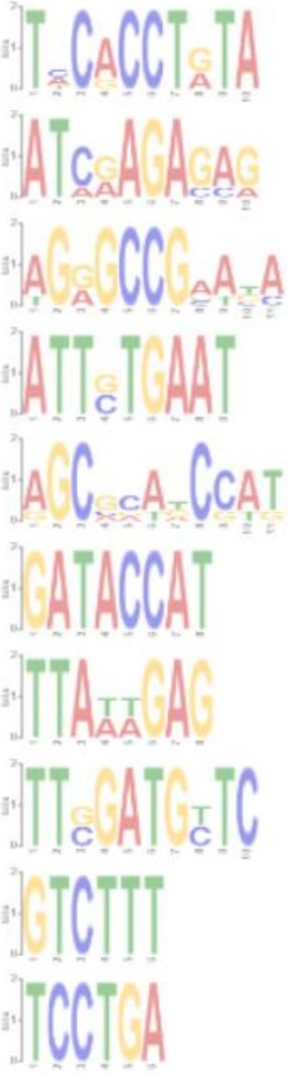

| | Logo | E-value | Sites | Width |
|---|---|---|---|---|
| 1. | | 2.1e+000 | 5 | 10 |
| 2. | | 1.7e+001 | 5 | 10 |
| 3. | | 1.1e+002 | 5 | 11 |
| 4. | | 5.7e+003 | 2 | 9 |
| 5. | | 6.3e+003 | 5 | 11 |
| 6. | | 7.2e+003 | 2 | 8 |
| 7. | | 1.6e+004 | 2 | 8 |
| 8. | | 1.2e+004 | 2 | 10 |
| 9. | | 1.7e+004 | 2 | 6 |
| 10. | | 2.4e+004 | 2 | 6 |

### DMCs

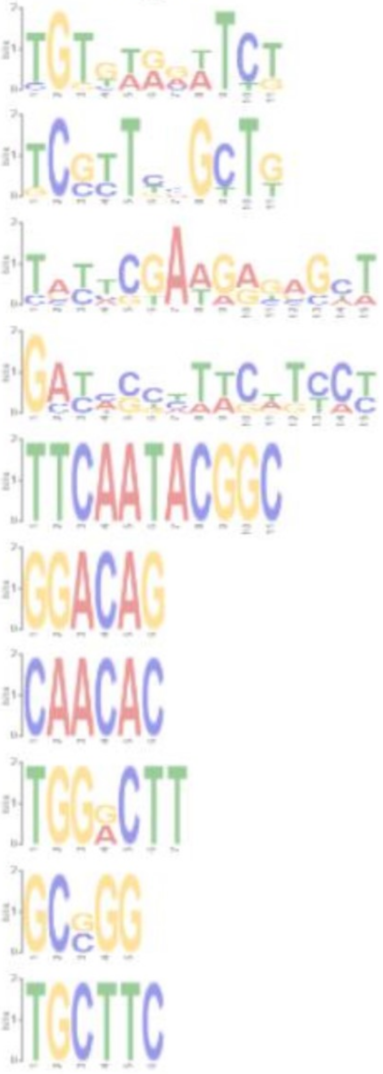

| | Logo | E-value | Sites | Width |
|---|---|---|---|---|
| 1. | | 8.6e+000 | 8 | 11 |
| 2. | | 6.1e+001 | 6 | 11 |
| 3. | | 2.1e+002 | 5 | 15 |
| 4. | | 1.5e+003 | 5 | 15 |
| 5. | | 1.9e+003 | 2 | 11 |
| 6. | | 1.3e+004 | 2 | 6 |
| 7. | | 2.1e+004 | 2 | 6 |
| 8. | | 3.2e+004 | 2 | 7 |
| 9. | | 1.9e+004 | 2 | 5 |
| 10. | | 2.1e+004 | 2 | 6 |

S5 Fig. MEME motif analysis results for the 49 and 47 DMAs and DMCs identified in the main analysis.15-nucleotide long sequences surrounding DMAs and DMCs were extracted, and motifs were analysed considering minimum and maximum motif widths of 4 and 15 nucleotides, respectively. A maximum number of 10 motifs were searched. No motifs reached significance (E-value < 0.05).

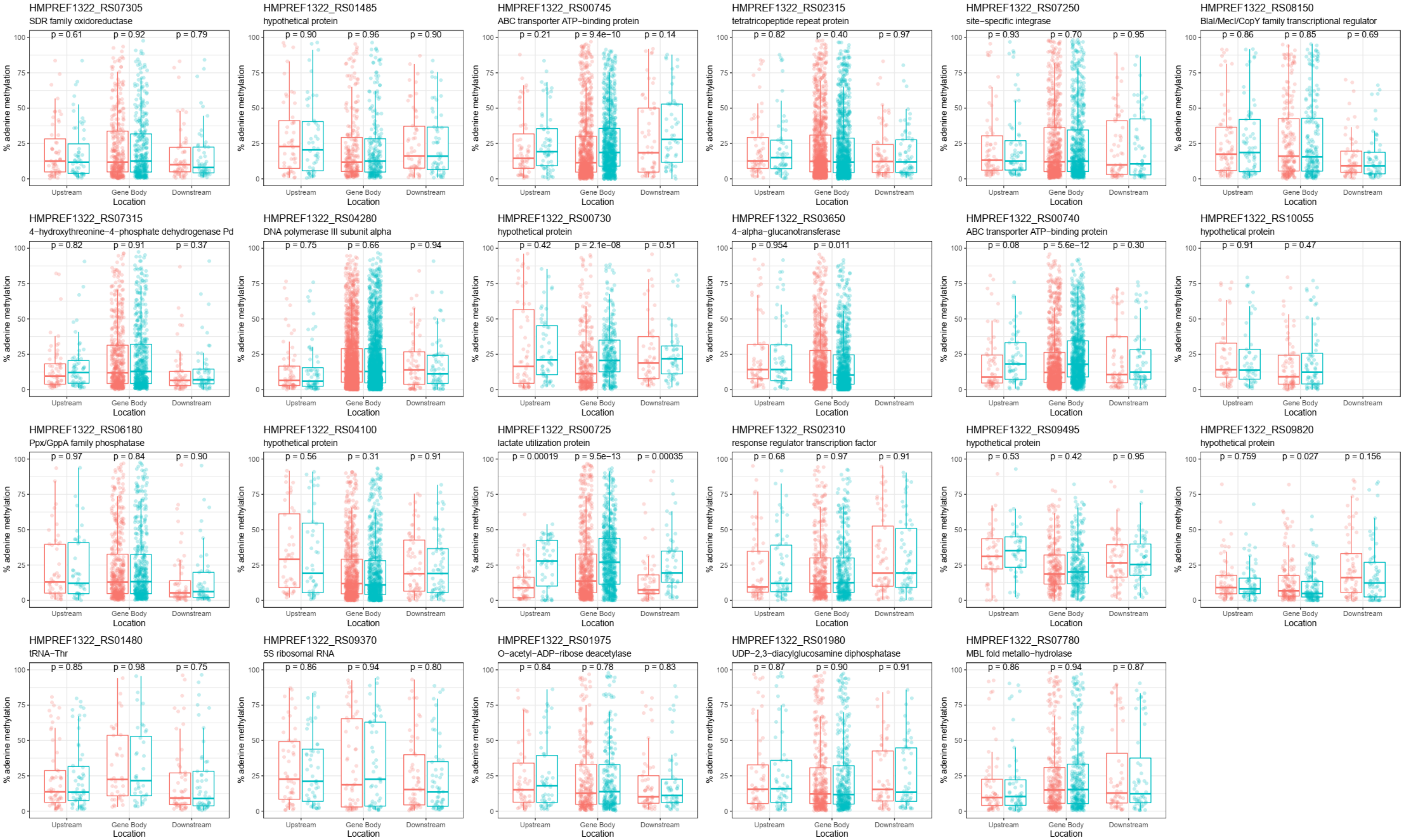


S6 Fig. Adenine methylation patterns upstream (-100 bp), downstream (+100 bp) and in the gene body of differentially expressed genes annotated to DMAs. Differential methylation significance was tested using Wilcoxon signed-rank tests between limited (red) and excess (blue) hemin conditions.

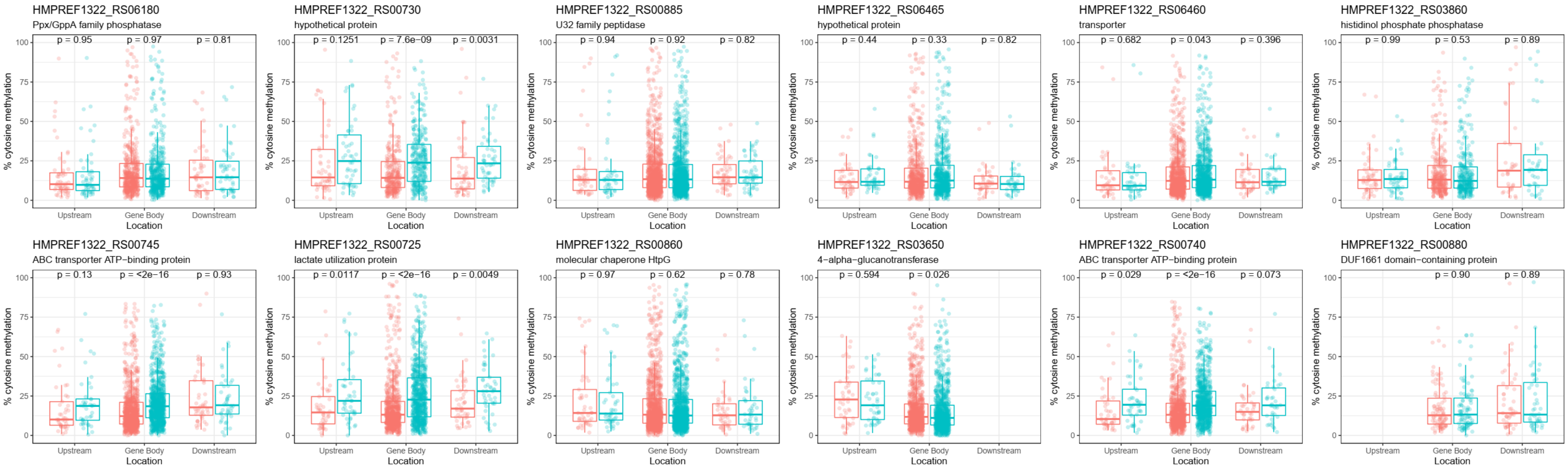


S7 Fig. Cytosine methylation patterns upstream (-100 bp), downstream (+100 bp) and in the gene body of differentially expressed genes annotated to DMAs. Differential methylation significance was tested using Wilcoxon signed-rank tests between limited (red) and excess (blue) hemin conditions.

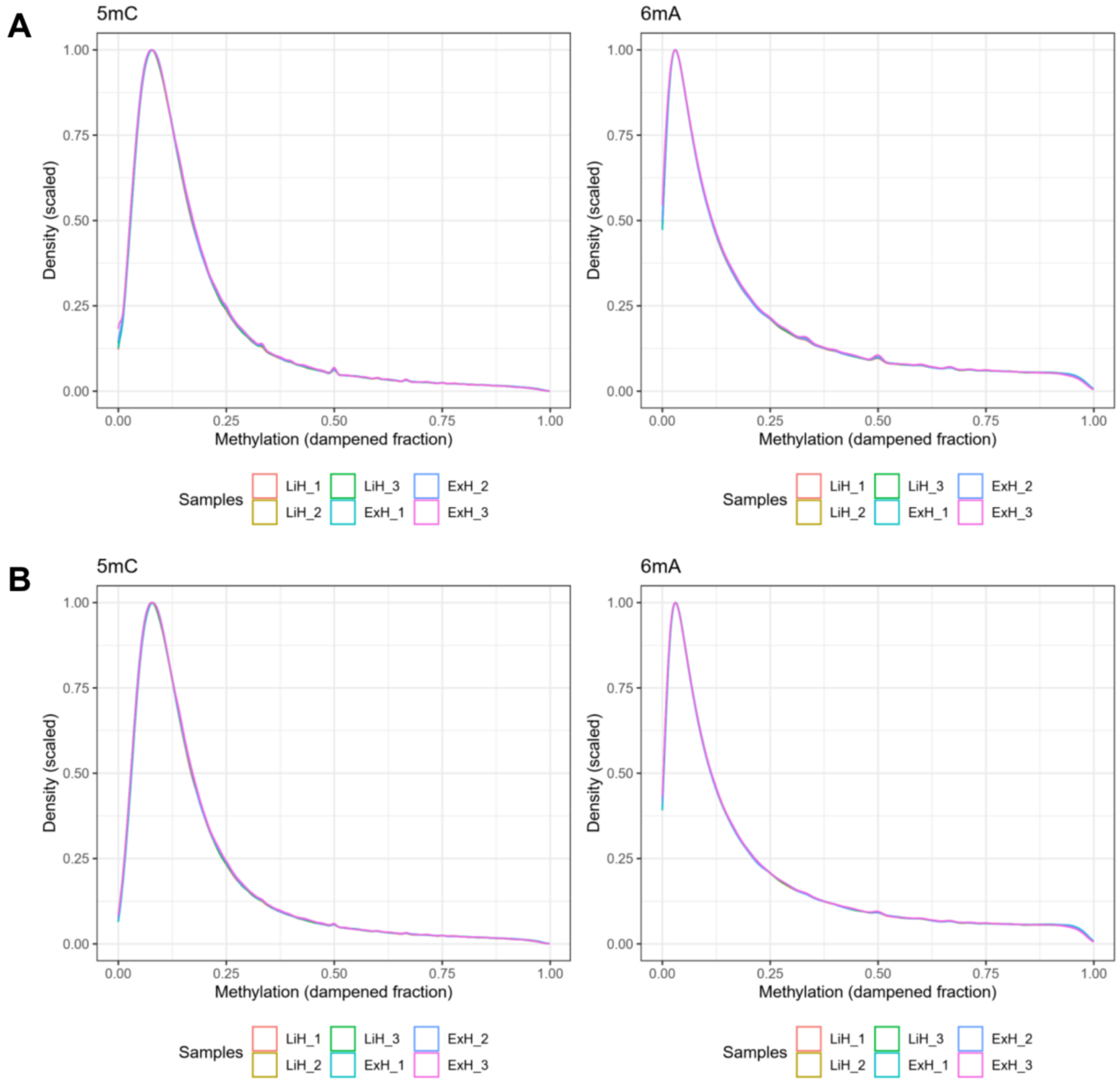


S8 Fig. Distribution of all-context DNA methylation for *P. gingivalis* growth in limited (LiH) and excess (ExH) hemin conditions, selecting for 10× (A) and 100× (B) coverage.

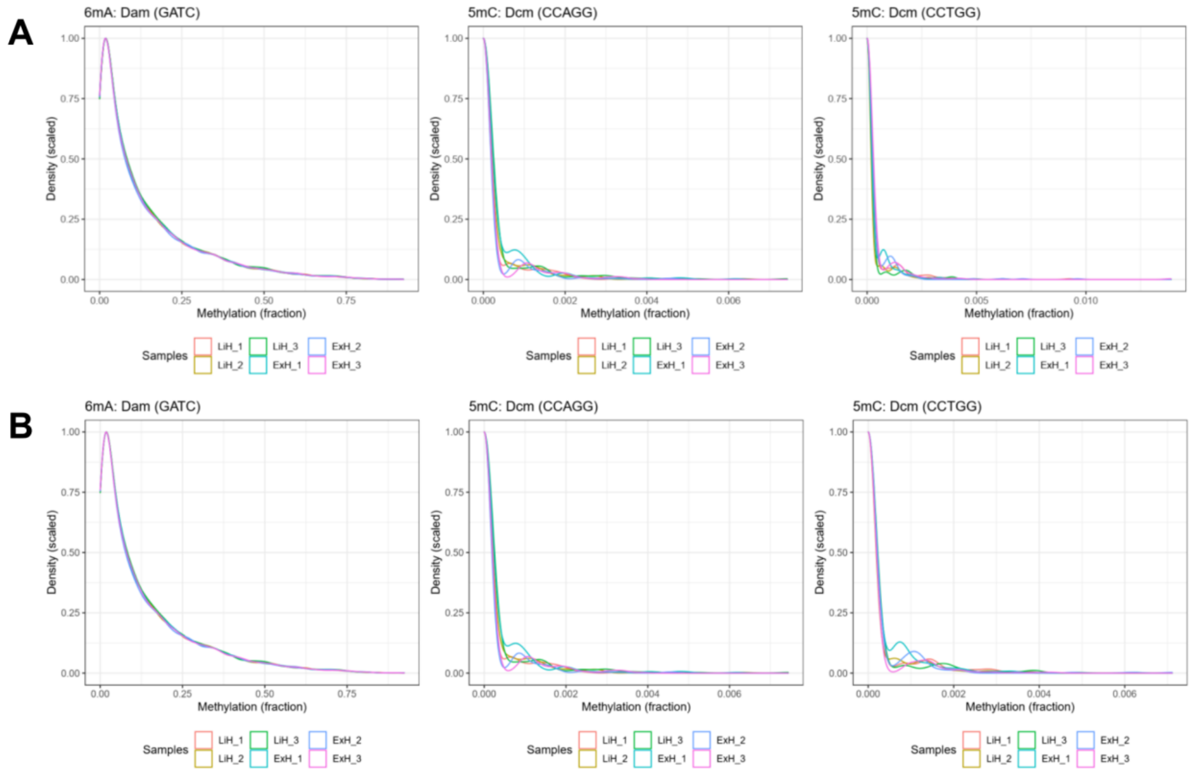


S9 Fig. Distribution of Dam/Dcm DNA methylation for *P. gingivalis* growth in limited (LiH) and excess (ExH) hemin conditions, selecting for 10× (A) and 100× (B) coverage.

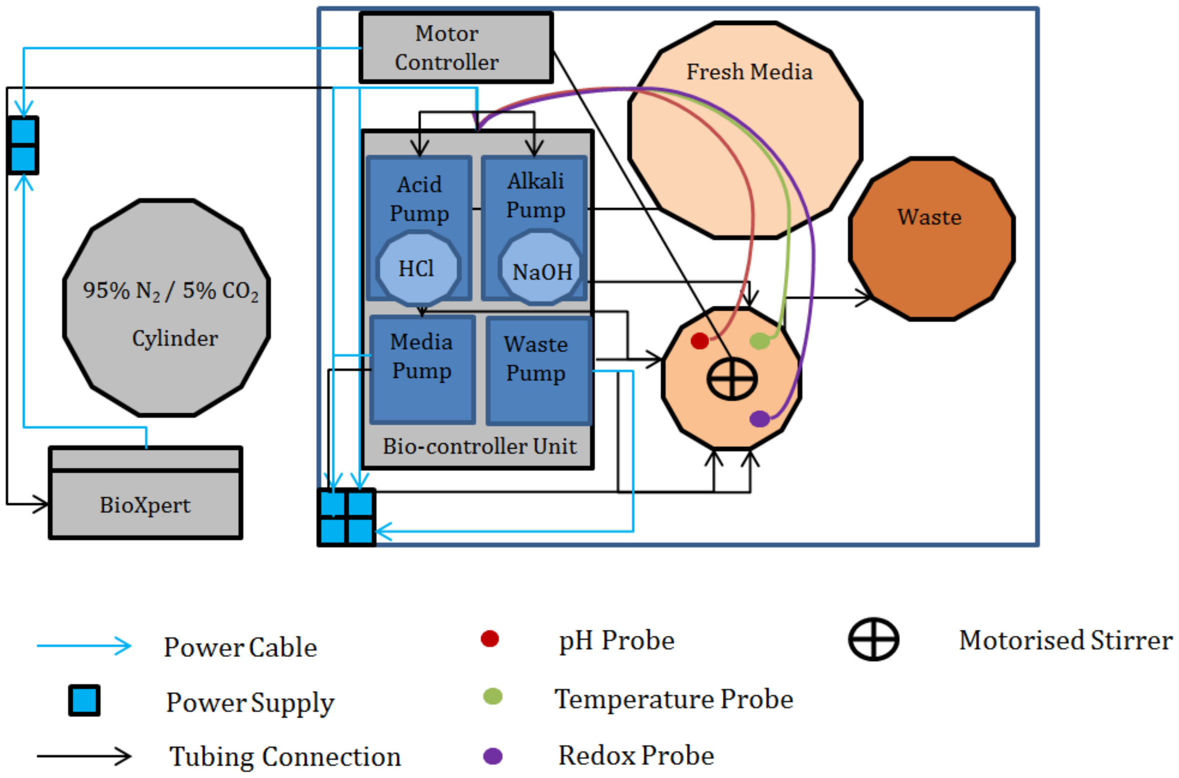


S10 Fig. Schematic representation of chemostat set up in class II fume cabinet.
